# Insights into the genomics of clownfish adaptive radiation: the genomic substrate of the diversification

**DOI:** 10.1101/2022.05.12.491701

**Authors:** Anna Marcionetti, Nicolas Salamin

## Abstract

Clownfishes are an iconic group of coral reef fishes that evolved a mutualistic interaction with sea anemones, which triggered the rapid diversification of the group. We investigated the genomic architecture underlying this process to determine the genomic characteristics associated with the adaptive radiation of the group and assess the mechanisms of parallel evolution in clownfishes.

We took advantage of the available genomic data of five pairs of closely related but ecologically divergent clownfish species to perform comparative genomic analyses. We found that clownfish genomes show two bursts of transposable elements, overall accelerated coding evolution, and topology inconsistencies potentially resulting from hybridization events. These characteristics possibly facilitated the rapid diversification of the group. We also detected a signature of positive selection throughout the radiation in 5.4 % of the clownfish genes. Among them, five presented functions associated with social behavior and ecology. They could have potentially played a role in the evolution of size-based hierarchical social structure so particular to clownfishes. Finally, we found genes with patterns of either relaxation or intensification of purifying selection and signals of positive selection linked with clownfish ecological divergence, suggesting some level of parallel evolution during the diversification of the group.

Altogether, these results provide the first insights into the genomic substrate of clownfish adaptive radiation. This work integrates the growing collection of studies investigating the genomic mechanisms governing species diversification, which brings us a step closer to understanding how biodiversity on Earth is created

## INTRODUCTION

Adaptive radiation is defined as the rapid diversification of an ancestral population into several ecologically different species, associated with adaptive morphological or physiological divergence (Schluter, 2000). This process is considered to play a central role in creating the spectacular diversity of life on Earth (Simpson, 1953; Schluter, 2000). For decades, researchers have been investigating the causes and consequences of adaptive radiations (e.g., Givnish & Sytsma, 1997; Schluter, 2000; Seehausen, 2004; Glor, 2012; Yoder et al., 2010; Givnish, 2015; Soulebeau et al., 2015; Stroud & Losos, 2016; Martin & Richards, 2019), with the ultimate goal to broaden our understanding of the mechanisms governing species diversification and, ultimately, the buildup of biodiversity.

A widespread phenomenon in adaptive radiations is convergent evolution, which describes the repeated evolution of similar phenotypes into distinct lineages (e.g., Schluter & Nagel, 1995; Rundle et al., 2000; Blackledge et al., 2004; Muschick et al., 2012; Vizueta et al., 2019). Convergent evolution is generally explained as the results of independent adaptation to similar ecological conditions (Schluter, 2000; Brakefield, 2006; Losos, 2011). While textbook examples of convergent evolution typically include the development of wings in birds and bats or the evolution of echolocation in bats and dolphins (Roberts, 1986; Vater & Kössl, 2004; Liu et al., 2010), in rapidly radiating lineages, convergent traits can also be observed between phylogenetically closer species. For instance, equivalent ecotypes of *Anolis* lizards (Losos, 2009) or *Tetragnatha* spiders (Blackledge & Gillespie, 2004) have arisen in the Greater Antilles and Hawaiian islands, respectively. Similarly, benthic and limnetic sticklebacks fish have evolved repeatedly in postglacial lakes (Schluter & Nagel, 1995; Rundle et al., 2000), and the occurrence of convergent forms of cichlids was observed both within the East African Lake Tanganyika (Muschick et al., 2012) and across the Tanganyika and the Malawi Lakes (Kocher et al., 1993). Thus, the adaptive radiation process provides an interesting setup to investigate not only the mechanisms behind species diversification, but also the repeatability and predictability of evolution, a central question in evolutionary biology (Rosenblum et al., 2014, Kingman et al., 2021).

With the advances in molecular genetics and sequencing technologies, the development, use, and availability of genomic tools for non-model systems have boomed (Abzhanov et al., 2008). Consequently, several studies have started investigating the intrinsic genomic factors that may promote adaptive radiations. Recent examples have included cichlids (Brawand et al., 2014; Faber-Hammond et al., 2019; McGee et al., 2020; Xiong et al., 2021), threespine sticklebacks (Jones et al., 2012; Verta & Jones, 2019), *Anolis* lizards (Feiner, 2016), Darwin’s finches (Lamichhaney et al., 2015), *Heliconius* butterflies (Dasmahapatra et al., 2012; Supple et al., 2013; Edelman et al., 2019) or *Dysdera* spiders (Vizueta et al., 2019). These studies found that a wide array of genome-wide changes, including chromosomal duplications, expansions of gene families, bursts of transposable elements (TEs), and accelerated evolution on coding and non-coding sequences, could predispose particular lineages to radiate adaptively (Jones et al., 2012; Brawand et al., 2014; Fan & Meyer, 2014; Feiner, 2016; Berner & Salzburger, 2015; Faber-Hammond et al., 2019; Verta & Jones, 2019; Xiong et al., 2021). Besides these overall genomic features potentially linked with rapid diversification, these studies provided evidence of the key role played by ancient polymorphism and hybridization events (both ancestral or between diverging lineages) in shaping adaptive radiations (e.g., Dasmahapatra et al., 2012; Berner & Salzburger, 2015; Lamichhaney et al., 2015; Meier et al., 2017; Malinsky et al., 2018; Edelman et al., 2019; Svardal et al., 2020; Kozak et al., 2021).

The question of whether convergent phenotypes - widely observed, for instance, in adaptive radiations - originated through shared genetic and molecular mechanisms (i.e., “parallel evolution” as defined in Rosenblum et al., 2014) has also started to be elucidated (Rosenblum et al., 2014; Elmer & Meyer, 2011; Sackton & Clark, 2019). Parallel evolution between different organisms was detected at diverse hierarchical levels, with, for instance, mutations at identical amino acid residues (Protas et al., 2006; Zhen et al., 2012; Projecto-Garcia et al., 2013), at distinct sites within the same locus (Kingsley et al., 2009; Rosenblum et al., 2010; Linnen et al., 2013) or in different genes within the same pathway (Arendt & Reznick, 2008). These cases arose from mutations that occurred independently in different species (see Table 1 in Stern, 2013). However, parallel evolution can also be achieved by the appearance of shared ancestral (and polymorphic) alleles - like in the convergent loss of lateral plates in the adaptive radiation of sticklebacks (Cresko et al., 2004; Colosimo et al., 2005) - or by introgressive hybridization - as in the convergent acquisition of wing patterns during the diversification of *Heliconius* butterflies (Dasmahapatra et al., 2012; Lewis et al., 2019).

**Table 1.** Positively selected genes in the whole clownfish clade showing particularly interesting functions. For each gene, the corresponding SwissProt ID and gene names are reported. Information on the log-likelihood of the null model (H0, no positive selection) and alternative model (H1, positive selection in clownfishes, Figure 1A) is reported. Likelihood-ratio test (LRT) multiple-testing corrected *q* values, the proportion of sites under positive selection on the tested branches (% PS sites) and the corresponding ω values are reported for each gene.

| OG ID | SwissProt ID | Protein Name | logL(H0) | logL(H1) | LRT $q$ Values | % PS sites | $\omega_2$ |
| --- | --- | --- | --- | --- | --- | --- | --- |
| OG20045_2b | Q90334 | Isotocin receptor (ITR) | -4555.44 | -4547.56 | 4.60E-003 | 0.3 | 51.142 |
| OG14285 | P01170 | Somatostatin-2 (SST2) | -1784.81 | -1768.73 | 4.31E-003 | 2.7 | 61.387 |
| OG16291 | Q9Y5X5 | Neuropeptide FF receptor 2 (NPFFR2) | -5198.93 | -5191.55 | 5.55E-003 | 1.0 | 10.922 |
| OG8036 | Q8HZK2 | Dual oxidase 2 (DUOX2) | -32014.46 | -32005.58 | 4.31E-003 | 0.4 | 14.285 |
| OG8544 | O18315 | Rhodopsin (RHO) | -4389.29 | -4374.97 | 1.12E-002 | 1.0 | 26.142 |

The study of adaptively radiating lineages has broadened our knowledge of the mechanisms behind species diversification. However, much remains to be understood on the overall genomic patterns associated with adaptive radiations, the mechanisms underlying the acquisition of convergent phenotypes, and the extent of parallel evolution. An interesting group to extend our understanding of these processes are clownfishes (or anemonefish). This iconic coral reef fishes (genera *Amphiprion* and *Premnas*) consists of 28 recognized species and two natural hybrids (Fautin & Allen, 1997; Ollerton et al., 2007; Gainsford et al., 2015). One distinctive characteristic of this group is the mutualistic interaction they maintain with sea anemones. Within the sea anemones, clownfishes live in a size-based social hierarchy (Fricke, 1979; Ochi, 1989; Buston, 2003) and are sequential hermaphrodites (Fricke & Fricke, 1977; Moyer & Nakazono, 1978; Fricke, 1979). While all species are associated with sea anemones, there is a large variability in host usage within the group. Some species are strictly specialists and can interact with a single species of sea anemones, while others are generalists and can inhabit up to ten hosts (Fautin & Allen, 1997; Ollerton et al., 2007; Gainsford et al., 2015). This mutualism acted as the key innovation that triggered the adaptive radiation of the group (Litsios et al., 2012). Following the acquisition of the interaction with sea anemones, the divergence in host usage likely drove the radiation, and within different clades, increasingly specialized species originated repeatedly and independently (Litsios et al., 2012). As a result, the diversifying species in different clades developed convergent phenotypes associated with host usage (i.e., generalists/specialists gradient; Litsios et al., 2012). While the primary radiation of the clownfishes happened in the Indo-Australian Archipelago, a second replicated and geographically independent radiation showing the same gradient of host usage occurred in the Western Indian Ocean (Litsios et al., 2014).

In this study, we investigated the genomic architecture of the clownfish adaptive radiation. Our first aim was to explore the genomic characteristics associated with the radiation of the clownfishes. In particular, we examined if bursts of TEs, increased gene duplications, signatures of accelerated evolution, and evidence of hybridization events - previously hypothesized to be associated with radiating lineages - were observed in clownfishes. We expected to detect at least some of these features, as they create the genomic variations necessary for natural selection to act, likely facilitating species diversification. We performed comparative genomic analyses of ten publicly available clownfish species covering the entire divergence and distribution range of the group (i.e., *P. biaculeatus, A*. ocellaris, *A. perideraion, A. akallopisos, A. polymnus, A. sebae, A. melanopus, A. bicinctus, A. nigripes*, and *A. frenatus*; Marcionetti et al., 2019). These species represent five pairs of closely related species showing ecological and phenotypic divergence within pairs, but ecological and phenotypic convergence between them; Litsios et al., 2012; see Figure 1 in Marcionetti et al., 2019).

**Figure 1.**
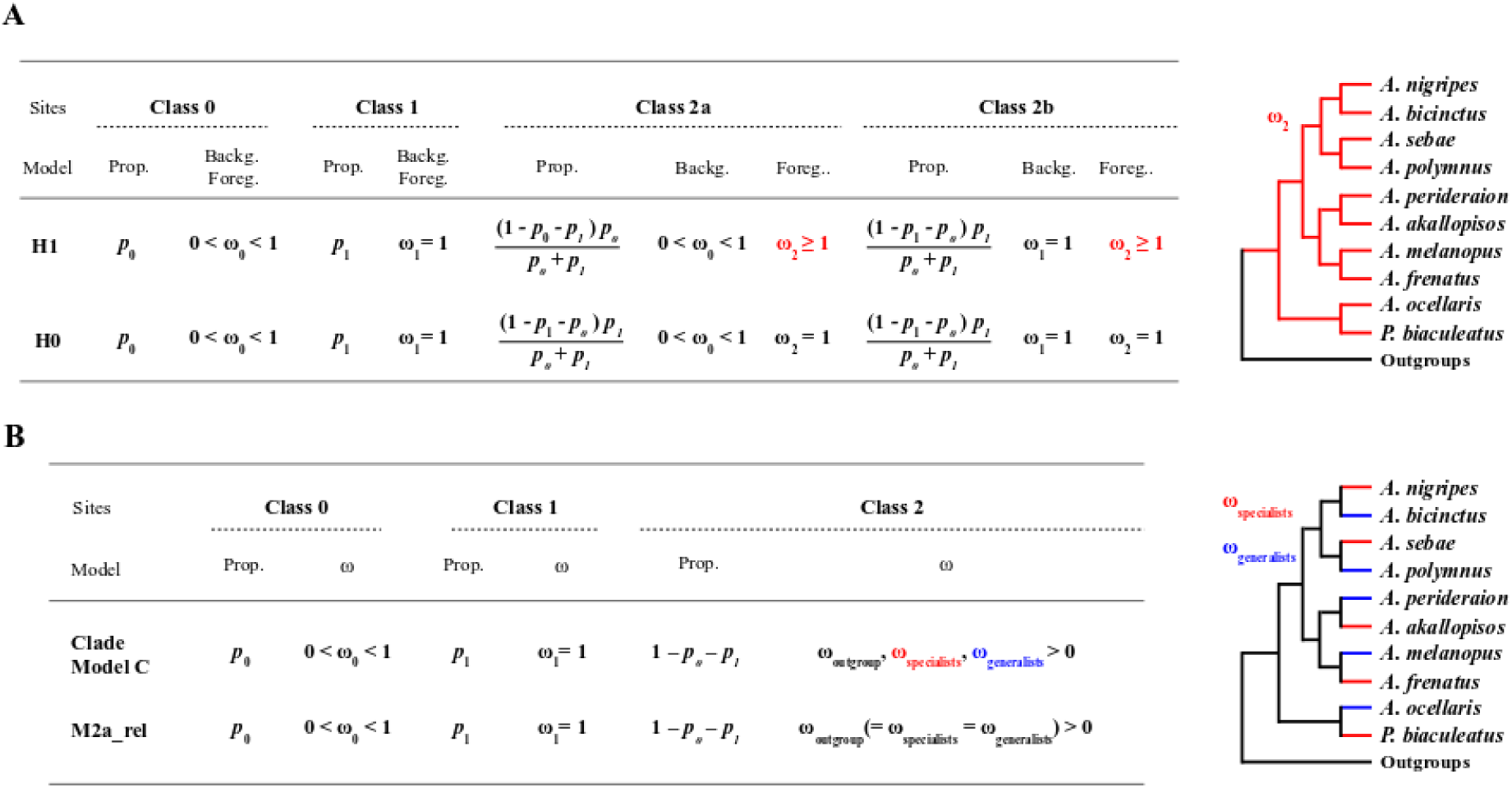
Models used for the positive selection analysis in the whole clownfish clade (A) and the analysis of selection linked with host and habitat divergence (B). A) In the branch-site model A, the null model with fixed ω_2_ (H0) was compared to the alternative model (H1), where ω_2_ was estimated. B) The null model M2a_rel with equal ω across branches was compared to the Clade Model C, where ω could vary in the three groups (i.e., specialists branches in red, generalists branches in blue and outgroups in black).

Our second aim was to determine the extent and the mechanisms of parallel evolution in the radiation of clownfishes. By taking advantage of these evolutionary replicates, we investigated whether the observed convergence at the phenotypic level was mirrored by convergent changes at the genetic level. The rationale behind this second aim was that genes involved in the ecological divergence of clownfishes might evolve at different rates or show different selective pressures associated with host usage. Suppose the same genes are involved in the ecological divergence of multiple species (i.e., parallel evolution). In that case, these genes should display similar evolutionary rates and selective pressures in all the concerned specialists and generalists species, or they might show topological inconsistencies in the case of adaptive introgressive hybridization (as in Dasmahapatra et al., 2012; Supple et al., 2013).

## MATERIAL AND METHODS

Whole-genome assemblies and annotations for the ten clownfish species (*Premnas biaculeatus, Amphiprion ocellaris, A. perideraion, A. akallopisos, A. polymnus, A. sebae, A. melanopus, A. bicinctus, A. nigripes*, and *A. frenatus*) and the lemon damselfish (*Pomacentrus moluccensis*) were taken from public repositories (Marcionetti et al., 2018: DRYAD Repository: https://doi.org/10.5061/dryad.nv1sv; Marcionetti et al., 2019: Zenodo Repository https://doi.org/10.5281/zenodo.2540241). We classified clownfish species depending on the number of interacting sea anemone species, resulting in either specialist (up to two sea anemones hosts) or generalist (more than two sea anemones hosts) species (Supplementary Table S1). Although the differential host and habitat use is more complex than this dichotomy, our classification separated clownfish species on the two principal axes of variation describing the mutualistic interaction and the morphological differentiation of the species(Litsios et al., 2012).

### Mitochondrial Genome Reconstruction

The mitochondrial genome of *A. frenatus* was available from Marcionetti et al. (2018; DRYAD Repository: https://doi.org/10.5061/dryad.nv1sv). We performed mitochondrial genome reconstruction of the nine additional clownfish species and the outgroup *P. moluccensis* as in Marcionetti et al. (2018). Briefly, we retrieved the sequenced reads from the SRA database (NCBI, BioProject ID: PRJNA515163). We randomly sub-sampled 20 million reads for each species and assembled them using MITObim (v.1.9; Hahn et al., 2013). We employed two different reconstruction methods, using either available mitochondrial genomes or barcode sequences to initiate the assembly. The NCBI accession IDs for the sequences used in both approaches are reported in Supplementary Table S2. We confirmed the consistency of the two reconstruction methods with Geneious (v.10.2.2; Kearse et al., 2012). We manually inferred the circularity of the sequences and we mapped the reads of the pool back onto the resulting mitochondrial genome to verify the reconstruction and assess the coverage using Geneious (v.10.2.2; Kearse et al., 2012).

### Orthology Inference between clownfishes and publicly available Actinopterygii genomes, codon alignments and gene tree reconstruction

Orthologous genes between the ten clownfish species, *P. moluccensis*, and 12 publicly available Actinopterygii species (*Astyanax mexicanus, Danio rerio, Gadus morhua, Gasterosteus aculeatus, Lepisosteus oculatus, Oreochromis niloticus, Oryzias latipes, Poecilia formosa, Takifugu rubripes, Tetraodon nigroviridis, Xiphophorus maculatus*, and *Stegastes partitus*) were obtained as reported in Marcionetti et al. (2019). We performed orthologous inference with OMA standalone (v.1.0.6, Altenhoff et al., 2013). We filtered the results to keep only hierarchical orthologous groups (HOGs) composed of both clownfish and outgroup species. We then classified HOGs as single-copy orthologous genes (1-to-1 OGs) or multicopy HOGs (Marcionetti et al. 2019). All HOGs were functionally annotated by appending the function of the genes composing them. The use of additional Actinopterygii species was necessary for positive selection analyses (see below) because the power to detect patterns of positive selection increases with increasing divergence and species (Anisimova et al., 2002). We also used the outgroup species to investigate the history of gene duplication in the clownfishes (see *Gene duplication analyses*).

We performed HOGs codon alignments as reported in Marcionetti et al. (2019). For each HOG, we produced protein alignments with MAFFT (v.7.130; Katoh & Standley, 2016), using the G-INS-i strategy and controlling for over-alignment with the *–allowshift* option. Codon alignments were obtained from protein alignments using PAL2NAL (Suyama et al., 2006). Because positive selection analyses are sensitive to alignment errors (Fletcher and Yang 2010), alignments were filtered to keep only high-confidence homologous regions following the approach done for the Selectome database (Moretti et al., 2014).

For each HOG, we reconstructed the gene tree from the unfiltered codon alignments with PhyML (v3.3; Guindon et al. 2010), applying both the HKY85 and GTR substitution models (100 bootstraps). We selected the best model with a likelihood ratio test (*df*=4). Unfiltered alignments were preferred to filtered alignments as the filtering steps frequently worsen single-gene phylogenetic inference (Tan et al., 2015).

### Mitochondrial and Nuclear Phylogenetic Trees

We investigated the relationship between the species by reconstructing mitochondrial and nuclear phylogenetic trees. We aligned the mitochondrial genomes of the ten clownfish species and the outgroup *P. moluccensis* with MAFFT (default parameters; v.7.450; Katoh & Standley, 2013). We visually checked the alignments to avoid poorly aligned regions and we reconstructed the mitochondrial phylogenetic tree with RaxML (v.8.2.12; Stamatakis, 2014) under the GTR+Г model, performing 100 bootstrap replicates. We reconstructed the nuclear phylogeny on the concatenated alignment of the 13,500 1-to-1 OGs. If missing genes were present in some species, gaps were introduced to maintain the correct concatenation. The nuclear phylogenetic tree was reconstructed with RaxML (v.8.2.12; Stamatakis, 2014) under the GTR+Г model (single model for the whole concatenated alignment), performing 100 bootstrap replicates. We did not use multispecies coalescent approaches because our goal was not to infer an accurate species tree but rather to investigate topological inconsistency in clownfishes (see below). Thus, it was only used as a reference topology.

We plotted the mitochondrial and nuclear phylogenies with the *cophylo* command of the R package phytools (v.0.6.44; Revell, 2012). The *P. moluccensis* individual was used to root the phylogenetic trees and was then removed from the plot.

### Topology inconsistency along the genome

We investigated the presence of topology inconsistencies reflecting potential hybridization events or incomplete lineage sorting across the nuclear genome of clownfishes. We considered scaffolds larger than 100 kb to reduce their number (2,508 scaffolds kept out of 17,801), while keeping most of the genomic information (80% of the total assembly length; Supplementary Table S3). In addition, we considered only scaffolds present in all clownfish species and *P. moluccensis*, the latter being used to root the phylogenetic trees.

Scaffolds were aligned using MAFFT (-auto parameter; v.7.130; Katoh & Standley, 2013). We filtered the resulting alignments with trimAl (--gappyout; v.1.4.1; Capella-Gutiérrez et al., 2009) to remove poorly aligned and gaps-rich regions. Alignment statistics before and after the filtering are available in Supplementary Table S3. We split the filtered alignments to obtain non-overlapping windows of 10 kb, 50 kb, or 100 kb. We only kept the last window of each alignment if its length was at least 60 % of the considered window length. We reconstructed phylogenetic trees for each window using PhyML (GTR+Г model, 100 bootstraps; v3.3; Guindon et al. 2010). We rooted the trees with *P. moluccensis*, which was removed before further analyses. We checked the support obtained for the trees by plotting the distribution of the bootstrap values of all nodes of all the trees and by calculating the average bootstrap support for each window (Supplementary Figure S1).

We visually investigated the tree topologies with DensiTree (v.2.2.5; Bouckaert, 2010), as implemented in the phangorn R package (v.2.4.0; Schliep, 2011). We summarized the topologies using the treespace R package (v.1.1.3; Jombart et al., 2017) by calculating the Robinson-Foulds distances (method=RF) before applying Metric Multidimensional Scaling (MDS). Groups of similar trees were identified by hierarchical clustering, using the function *findGroves* in the treespace R package (cutoff set to 380, see cluster dendrogram Figure 4A). Results obtained with the different window sizes (10 kb, 50 kb, and 100 kb) were consistent and we only considered windows of 100 kb. We visualized the MDS plot with ggplot2 (v.3.0.0; Wickham, 2016), and we plotted trees with the ggtree R package (v.1.14.6; Yu et al., 2017). We used the R package ape (v.5.2; Paradis and Schliep, 2019) to further explore the presence of topologies linked with host and habitat divergence (i.e. we searched for trees where specialist or generalist species branched together),

We mapped the scaffolds of *A. frenatus* against the chromosome-level assembly of *A. percula* (Lehmann et al., 2019; downloaded from Ensembl, Assembly AmpOce1.0, GCA_002776465.1) with blast (blastn v.2.7.1; https://blast.ncbi.nlm.nih.gov/Blast.cgi) to identify the genomic locations of the alternative topologies. We linked the scaffolds to the best chromosome hit and transferred the information of topological support for each window on the corresponding chromosome. We performed Gene Ontology (GO) enrichment analysis for specific regions of the genome (see below) using the topGO package (v.2.26.0; Alexa & Rahnenfuhrer, 2016) available in Bioconductor (http://www.bioconductor.org), setting a minimum node size of 10. Fisher’s exact tests with the weight01 algorithms were applied to examine the significance of enrichment, with p-values < 0.01 considered significant. We present here raw p-*values* instead of *p-*values corrected for multiple testing, following recommendations from the topGO manual.

### Transposable elements (TEs) annotation and analyses

We identified *de novo* transposable element (TE) families for the ten clownfish species and *P. moluccensis* with RepeatModeler (v.1.0.11; engine ncbi; Hubley & Smit, http://www.repeatmasker.org/RepeatModeler/), and we classified them with RepeatClassifier (within RepeatModeler) and TEClass (v.2.1.3, Abrusán et al., 2009). We complemented the obtained TE libraries with those of publicly available teleosts (*A. mexicanus, D. rerio, G. aculeatus, G. morhua, O. latipes, O. niloticus*, and *P. formosa*) downloaded from http://www.fishtedb.org/ (Shao et al., 2018). We annotated the TEs in clownfish and *P. moluccensis* genomes with RepeatMasker (v.4.0.7, http://www.repeatmasker.org/; Smit et al., 2015), using these TE libraries. We obtained information of TEs content in *A. percula* (Lehmann et al., 2019), *O. niloticus* (Brawand et al., 2014), *T. nigroviridis, G. aculeatus*, and *D. rerio* (Gao et al., 2016) for comparison with TEs content in clownfishes. We investigated the transposition history in clownfishes by performing a copy-divergence analysis of the TE superfamilies based on the Kimura 2-parameter distance (*K-*values; Kimura, 1980). We obtained the Kimura distance between each annotated TE copy and the consensus sequence of the respective TE family with the scripts calcDivergenceFromAlign.pl and createRepeatLandscape.pl provided in the RepeatMasker util directory.

### Gene duplication analyses

We investigated the gene duplication events occurring during the diversification of clownfishes and the other fish species *P. moluccensis, S. partitus, O. niloticus, G. aculeatus*, and *T. nigroviridis*. We retrieved 2,725 multicopy HOGs and filtered out the gene copies of the additional outgroup species not considered here. We employed a phylogenetic duplication analysis (PDA) approach (similar to Brawand et al., 2014), counting the number of gene duplication events observed at each branch of the phylogenetic trees of these species (see Supplementary Information S2 for details). The number of duplication events in each branch was then normalized to account for the divergence between species. We estimated the neutral genomic divergence between the species with RaxML (GTR+Г model, 100 bootstraps; v.8.2.12; Stamatakis, 2014), using ca. 7.5 million fourfold degenerate sites that we obtained from the codon alignments of 1-to-1 OGs. The number of duplication events detected on each branch was then divided by the corresponding branch length to obtain the duplication rate (i.e., the number of duplications normalized by the neutral divergence between the species; see Supplementary Information S2).

### Overall rate of evolution of clownfish genes

We explored the overall rate of evolution of clownfish genes by estimating the ratio of non-synonymous over synonymous substitutions (ω or Ka/Ks or dN/dS) in clownfish compared to the outgroups species. Values of ω smaller, equal, or larger than one correspond respectively to purifying selection, neutral evolution, and positive selection. With a homemade script, we randomly selected 20 1-to-1 OGs and concatenated their alignments. In the case of missing genes in some species, gaps were introduced to maintain the correct concatenation. We repeated the procedure 50 times to obtain 50 concatenated alignments of 20 randomly selected genes. For each alignment, we reconstructed the gene tree using PhyML (GTR model, 100 bootstraps v3.3; Guindon et al. 2010). We labeled the clownfish clade in the obtained trees and estimated the ω ratio in clownfishes and outgroups using the branch model implemented in *codeml* (PAML, v.4.9; Yang, 2007). We tested for a significant difference between the ω estimates obtained for the clownfishes and those obtained for the outgroups with a Welch Two Sample *t-*test.

### Positive selection analyses on the whole clownfish group

We tested for the presence of genes under positive selection in the whole clownfish clade. For single copy genes (1-to-1 OGs), we used the branch-site model implemented in *codeml* (PAML, v.4.9; Yang, 2007) on the filtered codon alignments and the species tree (see Supplementary Information S1). All branches of the clownfish group were set as foreground branches, while the remaining branches were assigned to the background (Figure 1A). In the null model, the foreground ω was constrained to be smaller or equal to 1, while in the alternative model, the foreground ω was estimated but was forced to be larger than 1 (Figure 1A). For each 1-to-1 OG, the best model was determined with a likelihood ratio test (LRT; *df*=1). We corrected the resulting *p-*values for multiple testing with the Benjamin-Hochberg method implemented in the *qvalue* package in R, following the approach used in the Selectome database (Moretti et al., 2013; false discovery rate (FDR) level of 0.1, the “robust” option, and the “bootstrap” method). To account for convergence issues encountered in the likelihood optimization in branch-site tests (Yang & Dos Reis, 2010), we fitted both the null and the alternative models three times.

We investigated the power and type I error in detecting positive selection by simulating data using *evolver* (PAML; v.4.9; Yang, 2007), following the approach used in Marcionetti et al. (2018). We simulated alignments of 1,000 codons under the branch-site model. We generated 20 trees following the species tree topology and with the branch lengths randomly drawn from the branch length distributions obtained from all the gene trees of the analyzed 1-to-1 OGs. For each tree, we simulated six sets of alignments with either no (ω_2_= 0.5 and ω_2_= 1) or an increasing level (ω_2_ of 2, 5, 10 and 20) of positive selection. We used the same pipeline as described above to estimate positive selection. We investigated the power to detect positive selection and the number of false positives (type I errors) by recording the number of significant LRT (*p-*value < 0.05) between the null model and the alternative model.

We retrieved the position on *A. frenatus* scaffolds of each gene under positive selection. This position was then transferred on the *A. percula* reference genome (see *Topology inconsistency along the genome*) to associate these genes to *A. percula* chromosomes. We selected the genes in the upper 90% of the foreground ω distribution and examined their function in the UniProt database (UniProt Consortium, 2018). We did gene ontology (GO) enrichment analysis by contrasting the GO annotation of all the significant genes against all analyzed genes (13,294 1-to-1 OGs) using the TopGO package (v.2.26.0; Alexa & Rahnenfuhrer, 2016) available in Bioconductor (http://www.bioconductor.org). Fisher’s exact tests with the weight01 algorithms were applied to examine the significance of enrichment, with p-values < 0.01 considered as significant. We present here *raw* p-*values* instead of *p-*values corrected for multiple testing, following recommendations from the TopGO manual. We extracted the genes associated with the significantly enriched GO terms and retrieved their functional information from the UniProt database (UniProt Consortium, 2018).

We investigated the presence of positive selection on some copies of the multicopy HOGs using the method aBSREL implemented in HyPhy (v.2.3.7; Smith et al., 2015). We performed the analysis in an exploratory way, testing for positive selection at each branch of the tree. Although this approach has a reduced power due to multiple testing, it was preferred as we did not know beforehand which copy of the genes could be positively selected. It however allowed us to obtain an overview of the selective pressures acting on the genes in all species. We corrected for multiple testing with the Benjamini-Hochberg method implemented in the *qvalue* package in R (FDR threshold of 0.1, “robust” option, and “bootstrap” method; Dabney et al., 2010). A similar analysis of multicopy genes was performed in Marcionetti et al. (2019), and we followed the same procedure here.

### Selection signatures associated with hosts and habitats divergence

We tested for signatures of selection associated with hosts and habitats divergence using the clade model C implemented in *codeml* (PAML v.4.9; Yang, 2007; Figure 1B), using the filtered codon alignments and the species tree (see Supplementary Information S1). Clade models allow differences in site-specific selective constraints among clades in the tree (Bielawski and Yang, 2004; Forsberg and Christiansen, 2003; Weadick and Chang, 2012). Here, we assigned clownfish species to either a specialist or generalist category, and we labeled the terminal branches of clownfishes with this host usage classification (Supplementary Table S1 and Figure 1B). The remaining branches were assigned to the background category. We fitted the clade model C, estimating separate ω ratios for each category in the analysis, and we compared it to the null model M2a_rel, where ω is fixed among clades (Figure 1B), with a LRT (*df*=2). The resulting *p-*values were corrected for multiple testing with the Benjamini-Hochberg method implemented in the *qvalue* R package (FDR threshold of 0.1, “robust” option, and “bootstrap” method; Dabney et al., 2010). To account for convergence issues potentially encountered in the likelihood optimization, we ran both models five times and kept the run resulting in the best likelihood.

Significant genes (q-value < 0.01) were classified into two main groups: OGs with patterns of purifying selection in both clownfish groups (ω_specialists_ and ω_generalists_ < 1) and OGs with signatures of positive selection in all specialists and/or generalist species (ω _specialists_ or/and ω generalists > 1).

For the first group, we looked for genes showing a different pattern of purifying selection in either specialists or generalists species. For this, we estimated the distribution of ω_background_, ω_specialists_ and ω_generalists_ and tagged genes falling in the lower 10% of the ω_specialists_ distribution but in the upper 10% of the ω_generalists_ and ω_background_ distributions as those experiencing intensified purifying selection in specialists (Figure 2A). Likewise, genes experiencing intensified purifying selection in generalists were obtained by considering the genes in the lower 10% of the ω_generalists_ distribution and in the upper 10% of the ω_specialists_ and ω_background_ distributions (Figure 2B). Finally, genes experiencing relaxed purifying selection in either specialists or generalists species were retrieved by considering those genes falling in the upper 10% of respectively the ω_specialists_ or ω_generalists_ distributions, but in the lower 10% of the remaining ω distributions (Figure 2C-D) Analyzing these results by considering the estimated ω_background_ of each gene was necessary to differentiate between intensified purifying selection in one group and relaxed purifying selection in the other.

**Figure 2.**
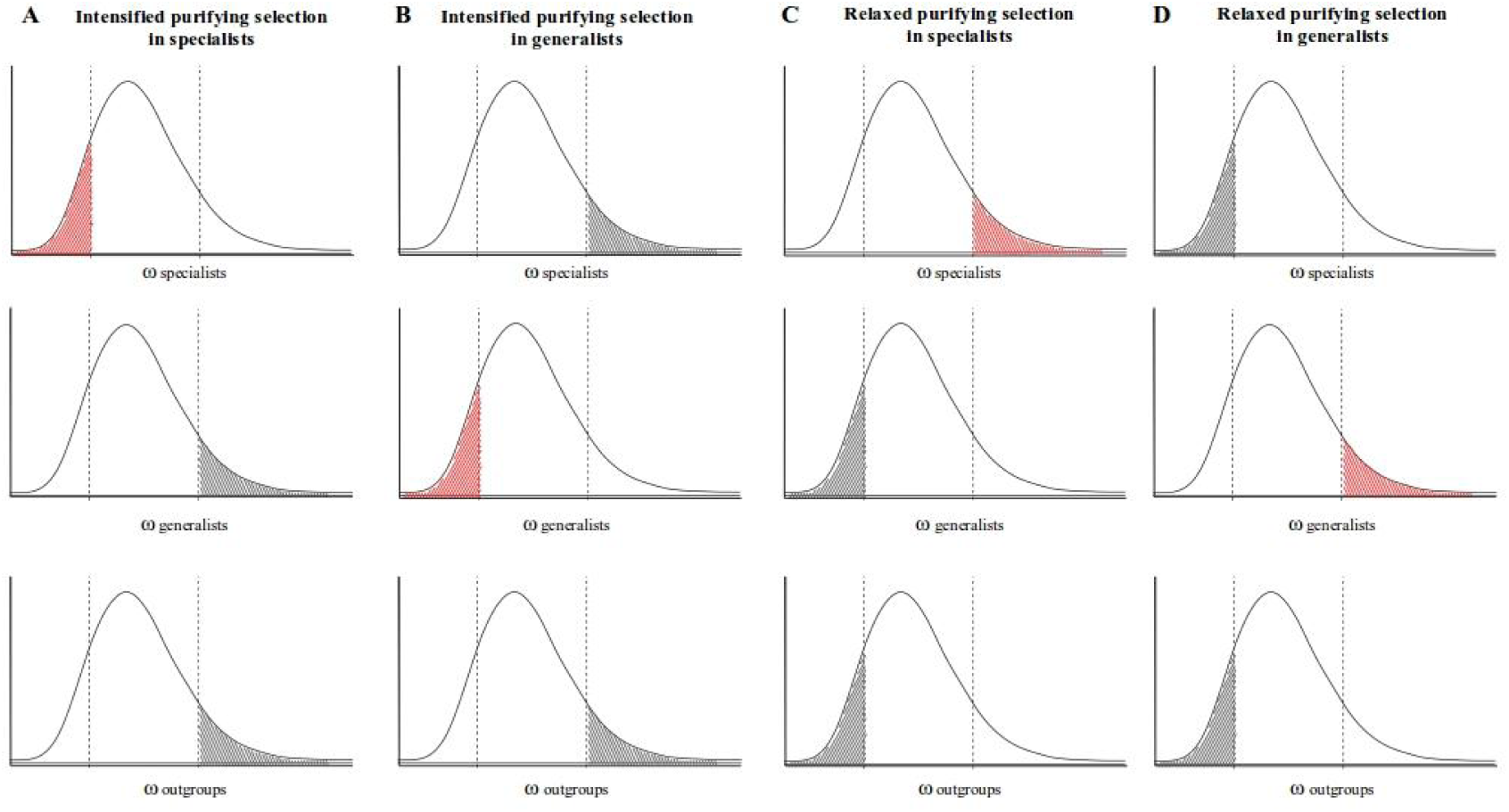
Illustration of the method used to define the categories of genes showing intensification or relaxation from purifying selection. Plots represent the distribution of ω estimates in specialists, generalists, and outgroups. Dotted lines represent the lower and upper 10% of the distribution. We defined genes with intensification of purifying selection in specialists (A) and generalists (B) and genes with relaxed purifying selection in specialists (C) and generalists (D).

For the second category, significant results were classified into genes positively-selected only in specialists (ω_specialists_ > 1.5; ω_generalists_ and ω_background_ <=1) and genes positively-selected only in generalists (ω_generalists_ > 1.5; ω_specialists_ and ω_background_ <=1). We selected ω > 1.5 as the threshold (instead of simply > 1) to exclude genes with a weak signal of selection and avoid false positives.

We manually investigated the functional annotation of the genes identified with the procedure described above by performing GO enrichment analyses with the TopGO package (v.2.26.0; Alexa & Rahnenfuhrer, 2016) available in Bioconductor (http://www.bioconductor.org). We contrasted the GO annotation of the significant genes in each category against all analyzed genes (13,294 1-to-1 OGs) and followed the same procedure as for the positively selected genes in the whole clownfish group.

### Evolutionary rate linked with host and habitat usage

We retrieved the reconstructed gene trees for each 1-to-1 OG (see Supplementary Information S1), and we considered the branch length as a proxy of the evolutionary rate of the OGs. We investigated the presence of OGs with the evolutionary rate linked with host and habitat usage by calculating the difference in branch length between generalists and specialists in each pair of closely related species. We considered the five pairs as replicates, and for each OG, we tested whether the difference was significantly different from zero with a one-sample Student t test. We corrected for multiple testing using the *p*.*adjust* function in R (method FDR). Additionally, for each pair of closely related species, we computed the distribution of the difference in branch length between generalists and specialists. The genes in the lower and upper 5% of the distribution were defined as those with a higher evolutionary rate in specialists and generalists, respectively (see Figure 7A). We investigated the intersection of these genes in the five pairs of closely related species to obtain genes that have a parallel increase of evolutionary rate in either all specialist or generalist species. We plotted the results with the function *venn* (venn R package, v. 1.10; Dusa, 2021). We tested whether the number of genes shared between 2, 3, 4 and 5 species pairs was significantly different between generalists and specialists by performing two-sample Student t tests for each category.

## RESULTS

Information on the nuclear genome assembly and annotation for the ten clownfish species and *P. moluccensis* is available in Marcionetti et al. (2019). Details on the hierarchical orthologous groups (HOGs) between clownfishes and the 13 Actinopterygii outgroup are also reported in Marcionetti et al. (2019). We obtained a final set of 15,940 HOGs that contained 13,500 single-copy orthologous genes (1-to-1 OGs). The remaining HOGs were classified as multicopy orthologs. We used the genome assemblies and this set of genes to investigate the genomic architecture and the molecular evolution associated with the adaptive radiation of clownfishes and to test for patterns of parallel evolution in specialists and generalists species.

### Mosaic genomes in clownfishes

We investigated the presence of topological inconsistencies between the mitochondrial and the nuclear genomes, as well as throughout the nuclear genome of clownfishes. Cytonuclear incongruences and topological disparities suggest that hybridizations or incomplete lineage sorting (ILS) occurred in the group. Additionally, inconsistencies where ecologically similar species branch together could indicate introgressive hybridization potentially involved in the buildup of the convergent phenotypes.

In both mitochondrial and nuclear phylogenetic trees, each species branched with the expected closely related pair (Figure. 3), which confirmed that the ten species formed five pairs of phylogenetically close but ecologically distinct species. Nevertheless, we found cytonuclear discordance at deeper nodes in the tree. Indeed, the pairs *A. melanopus - A. frenatus* and *A. perideraion - A. akallopisos* formed two sister groups in the nuclear phylogeny, while the latter was basal to *A. melanopus – A. nigripes* in the mitochondrial phylogenetic tree (Figure 3). This cytonuclear discordance was highly supported, with bootstrap support for the node higher than 0.95 (Figure 3).

**Figure 3:**
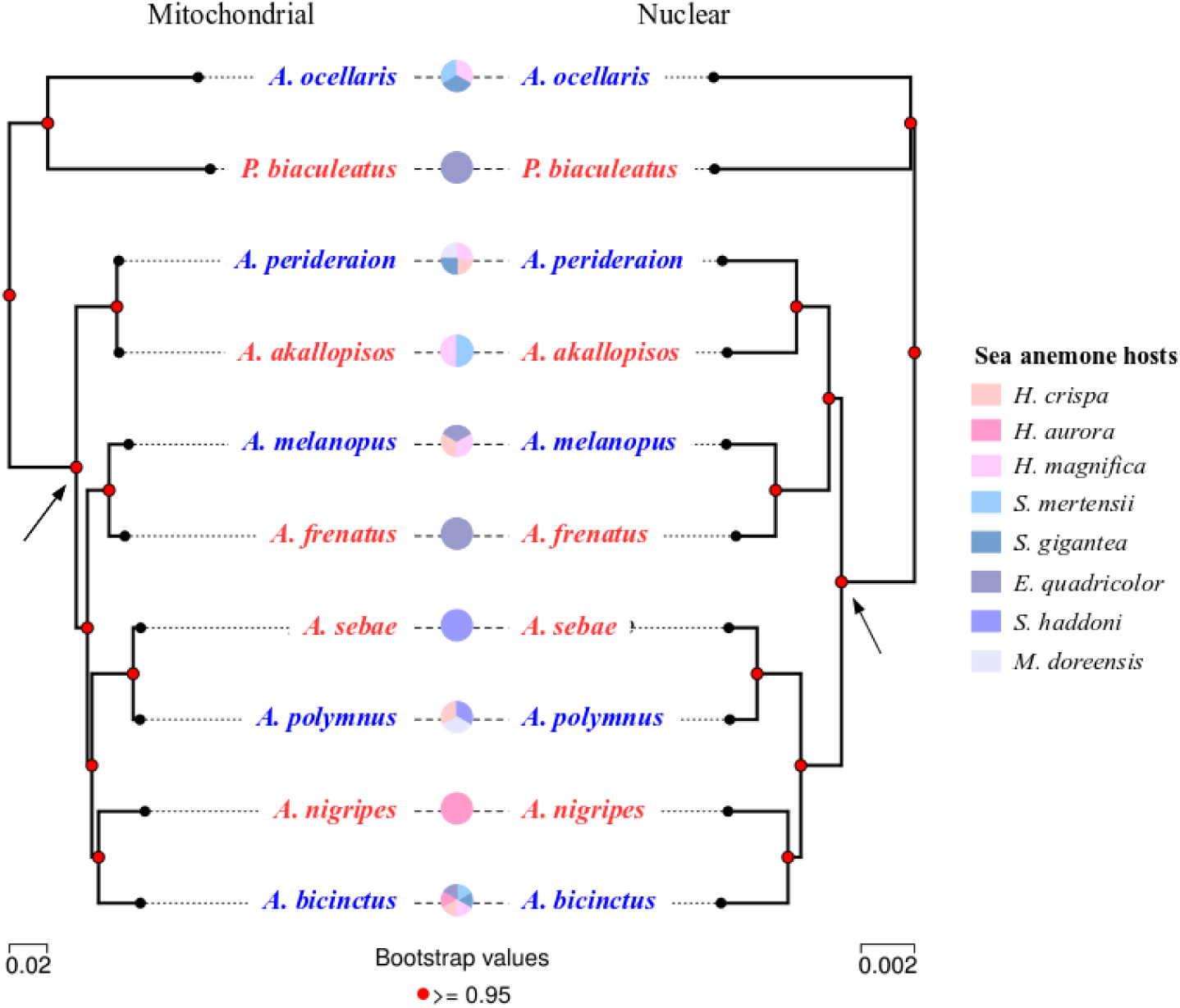
Mitochondrial and nuclear phylogenetic trees of the ten clownfish species. The mitochondrial phylogenetic tree was obtained with RAxML from the whole mitochondrial genome alignments (total of 16,747 bp). The nuclear phylogenetic tree was obtained with RaxML from the concatenated alignment of 13,500 orthologous genes (total of 22.5 Mbp). For both trees, 100 bootstrap replicates were performed. All nodes have bootstrap support higher than 0.95. The black arrows illustrate the discordance between the two topologies. The possible interactions between clownfish and sea anemones species are shown, with generalist and specialist species reported in blue and red, respectively.

For the analysis of topological inconsistencies along the genome, phylogenetic reconstruction for non-overlapping windows of 100 kb resulted in a total of 5,936 trees. Overall, the trees were well supported, with an average bootstrap value higher than 0.95 for 78 % of the windows (99.7 % when considering average bootstrap support higher than 0.8; Supplementary Figure S1).

Visual investigation of the tree topologies showed the presence of topological disparities in deep nodes, while the five pairs of closely related species predominantly branched together (Supplementary Figure S2). We quantified the different topologies along the nuclear genome and found five major clusters of trees (Figure 4). Clusters 1 and 2 contained 27% and 57% of the 5,936 gene trees, while the three others included far fewer (5%, 3%, and 2% of the trees for clusters 3, 4, and 5, respectively). The remaining trees (6%) showed a larger number of topologies, and thus we did not consider them as a cluster (Figure 4B, Supplementary Figure S3). All clusters contained trees with two different topologies that differed only by the branching of *P. biaculeatus* (depicted in Figure 4C by dotted lines). A slight majority of trees in each cluster placed *P. biaculeatus* as the sister species of *A. ocellaris* (between 53% and 74% of the gene trees), except in cluster 5, where *P. biaculeatus* was found at this position 90.8% of the time (Figure 4C).

**Figure 4:**
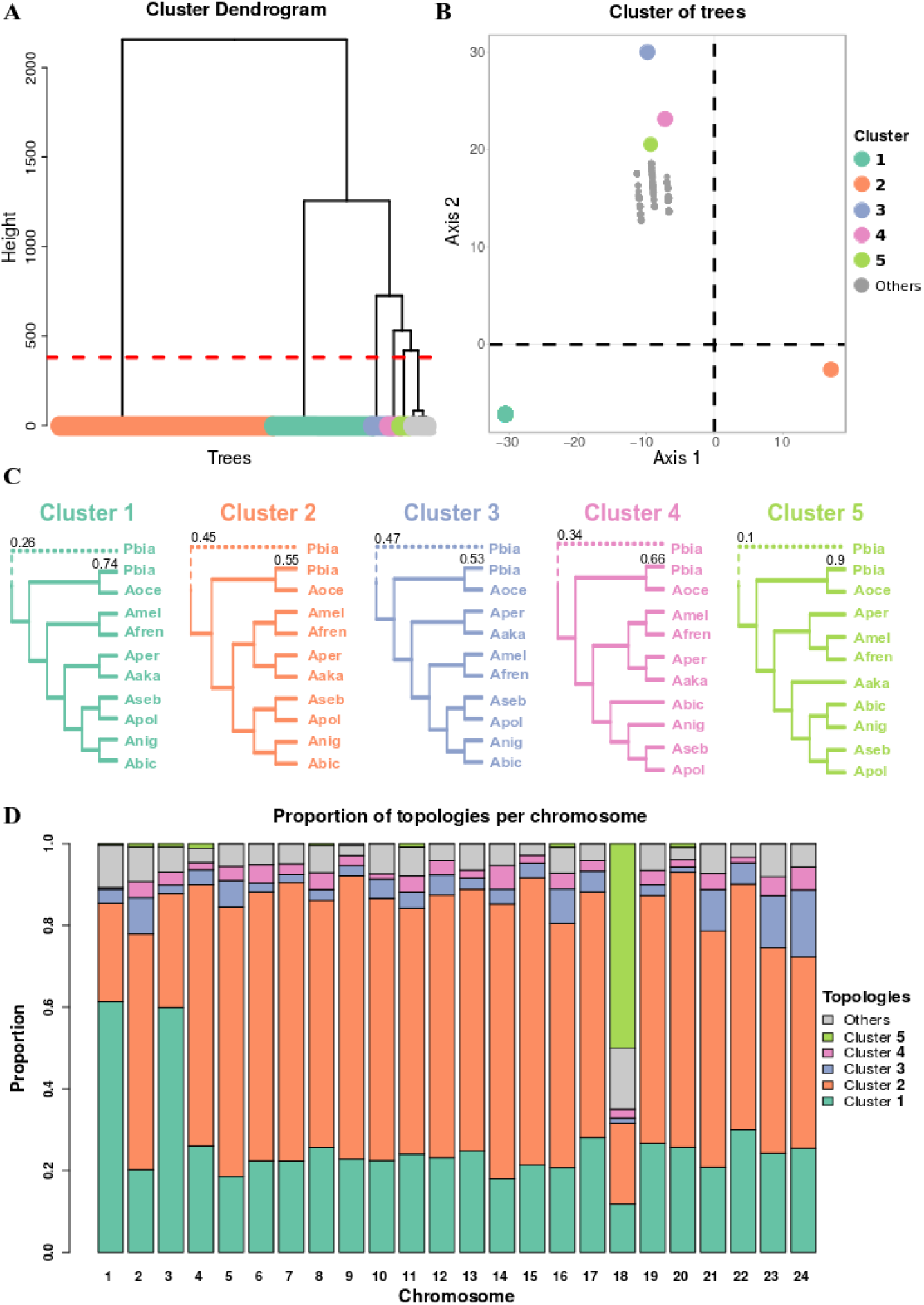
The five major topologies observed along the nuclear genome of clownfishes. A) Hierarchical clustering based on the Robinson-Foulds distance between the trees reconstructed along the genome. The red line corresponds to the cutoff set to define the five major clusters of trees. The last cluster (gray) is not considered because it is composed of different topologies (see DensiTree plot in Supplementary Figure S3). B) Multidimensional scaling plot of the five clusters of trees. C) The topologies of the five main clusters of trees represented in the genome. Pbia, Aoce, Amel, Afre, Aper, Aaka, Abic, Anig, Aseb, and Apol correspond to *P. biaculeatus, A. ocellaris, A. melanopus, A. frenatus, A. perideraion, A. akallopisos, A. bicinctus, A. nigripes, A. sebae*, and *A. polymnus*, respectively. The position of *P. biaculeatus* in the trees forming each cluster was variable, and the numbers on the top of the branches represent the proportion of trees showing the corresponding topology. D) Distribution of the five topologies across the chromosomes of *A. percula*. The additional topologies, reported in gray, are variable (see DensiTree plot in Supplementary Figure S3).

The main differences between the topologies of these five clusters were given by the branching of the *A. akallopisos – A. perideraion* pair. Indeed, *A. akallopisos – A. perideraion* and *A. melanopus – A. frenatus* were sister groups in cluster 2. This topology was the most represented in the genome and corresponded to the nuclear phylogenetic tree (Figure 3) and the expected species topology. In cluster 1, the pair *A. akallopisos – A. perideraion* was basal to the *A. bicinctus – A. nigripes – A. polymnus – A. sebae* complex (Figure 4C). Cluster 3 was characterized by the *A. akallopisos – A. perideraion* pair being basal to the *A. melanopus – A. frenatus* pair, which is the topology of the mitochondrial phylogenetic tree (Figure 3). In cluster 5, *A. perideraion* and *A. akallopisos* were not branching as sister species, but *A. akallopisos* was basal to the *A. bicinctus – A. nigripes – A. polymnus – A. sebae* group, while *A. perideraion* was basal to the *A. frenatus – A. melanopus* pair (Figure 4C). Cluster 4 was similar to cluster 2, but with *A. bicinctus* being basal to *A. nigripes* (Figure 4C).

The five topologies were distributed across the genome, with 1,053, 1774, 224, 165, and 55 scaffolds showing topologies of clusters 1 to 5, respectively (Figure 4D). The pattern was similar for most of the 24 chromosomes, which had the expected species tree (cluster 2) as the most frequently observed topology. Nevertheless, we observed three exceptions on chromosomes 1, 3, and 18 (Figure 4D). On chromosomes 1 and 3, the most abundant topology (around 60% of the windows) was cluster 1 (Figure 4D). Chromosome 18 showed a completely different scenario, with the topology of cluster 5 being the most frequent. Indeed, 88% of the windows supporting this topology were located on chromosome 18, and, altogether, they accounted for 50% of the topologies observed on this chromosome (Figure 4D). In these regions of chromosome 18, we identified 432 genes, of which 331 were functionally annotated. Gene Ontology (GO) enrichment analysis of these genes resulted in 24 enriched terms (*p-*value < 0.01; Supplementary Table S4) associated with the morphogenesis of the epithelium (GO:1905332), fertilization (GO:0009566), axis elongation (GO:0003401), and retina vasculature development (GO:0061298).

None of the five clusters of trees was summarized by topologies with specialists or generalists species branching together. Nonetheless, when we considered the remaining and more variable trees found in the sixth cluster (represented in gray in Figure 4A, B, D), we observed 13 topologies in which the specialist species *A. nigripes* and *A. sebae* branched as sister species (Supplementary Figure S4) with a bootstrap support of 100 %. The windows showing these topologies were located on ten different chromosomes and contained 57 annotated genes (Supplementary Table S5). Among them, we found the genes encoding for olfactory receptors (*OR52N5*, Olfactory receptor 4S1; *OR4S1*, Olfactory receptor 4S1) and the gene *Has2*, encoding for the hyaluronan synthase 2.

### Transposable elements content

We investigated transposable elements (TEs) content and transposition history in clownfishes to determine if TEs potentially played a role in the diversification of the group. We found that approximately 23-25% of clownfish genomes consisted of TEs, with TEs proportion and composition similar to what observed in *A. percula* and *O. niloticus* genomes (Supplementary Figure S5 and Supplementary Table S6). We detected two major TE bursts in the clownfish genomes (Kimura distance *K-value* of 0.05-0.06 and 0.18-0.19; Figure 5 and Supplementary Figure S6). The most recent burst was also present in *P. moluccensis*, although it was less pronounced. Indeed, at *K-value* of 0.05 and 0.06, the average TEs percent in clownfish genome was respectively 1.07% (SD=0.06) and 1.14% (SD=0.05), while in *P. moluccensis*, TEs represented only 0.45% and 0.53% of the genome, respectively (Supplementary Figure S6). When comparing the Kimura distance to the neutral genomic divergence between clownfish and the outgroup (see species divergence, Supplementary Figure S7), we found that the older TE burst (*K-value* 0.18-0.21) happened at the time of the split of *O. niloticus* with the common ancestor of the Pomacentridae. The more recent burst (*K-value* of 0.05-0.06) was situated around the split of *P. moluccensis* from the common ancestor of clownfishes.

**Figure 5.**
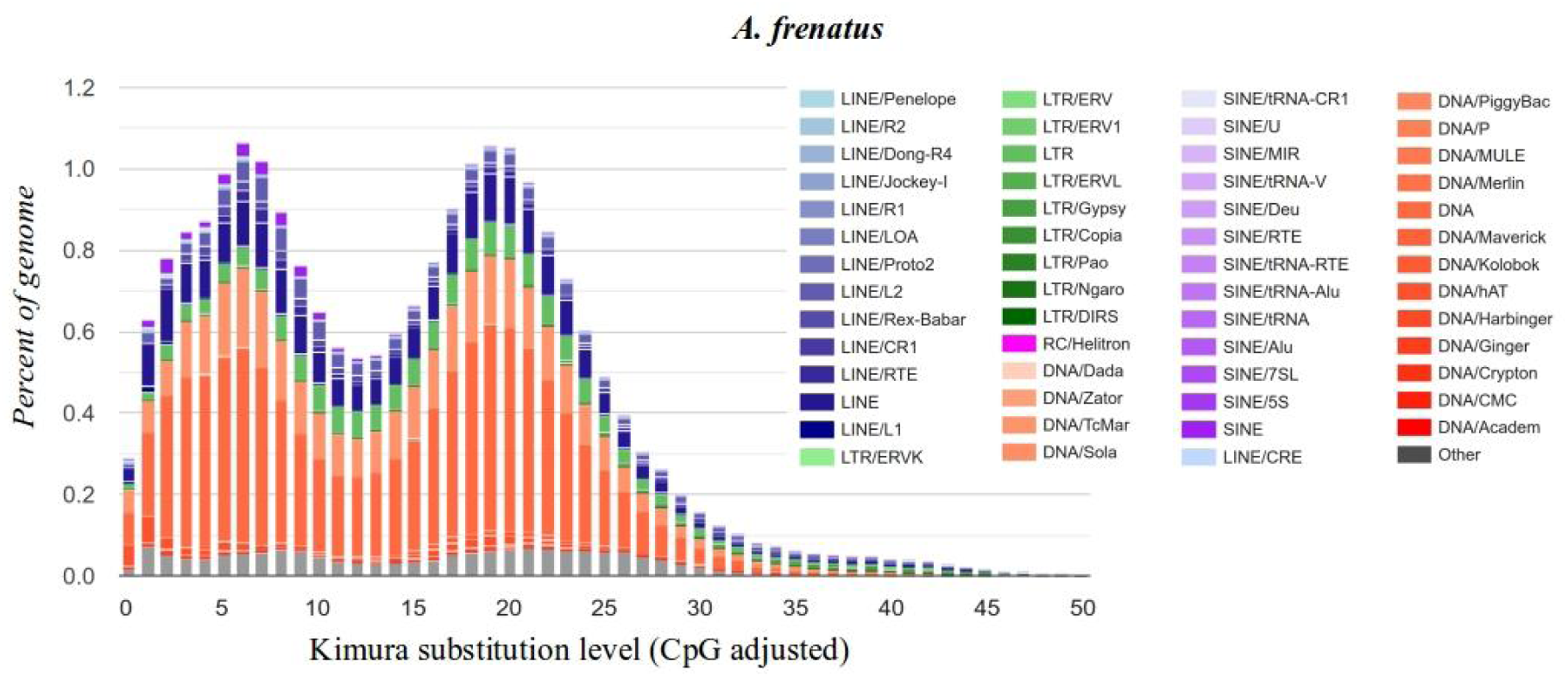
Kimura distance-based copy divergence analysis of transposable elements in *A. frenatus*. The different colors represent the different TE superfamilies. TEs were clustered according to Kimura distances to their corresponding consensus sequence. Copies clustering on the left side of the graph did not greatly diverge from the consensus sequence and potentially corresponded to recent events, while sequences on the right side likely corresponded to older divergence. Peaks in the graph indicate TE bursts. Similar results were obtained for the additional nine clownfish (Supplementary Figure S6).

### Gene duplication rate and positive selection on multicopy genes

We explored whether clownfishes showed an increased rate of gene duplication that could be associated with their diversification. We found that the duplication rate in the common ancestor of clownfishes was similar to the one observed for the *P. moluccensis -* clownfishes split and to the one in the common ancestor of the Pomacentridae (around 12 duplicated genes/percent of divergence; Supplementary Information S2 and Supplementary Figures S7). Additionally, we did not detect any multicopy gene with evidence of positive selection throughout the whole clownfish clade, and thus potentially associated with the group’s diversification (Supplementary Information S2).

### Overall accelerated evolution and positively selected single-copy genes in clownfish

We investigated the rate of protein-coding sequence evolution in clownfishes and found a significantly higher ω value in clownfishes (M=0.14, SD=0.03) compared to the outgroups (M=0.09, SD=0.02; t(73)=11.0, *p* < .001; Figure 6), which suggests an overall accelerated evolution in this group.

**Figure 6:**
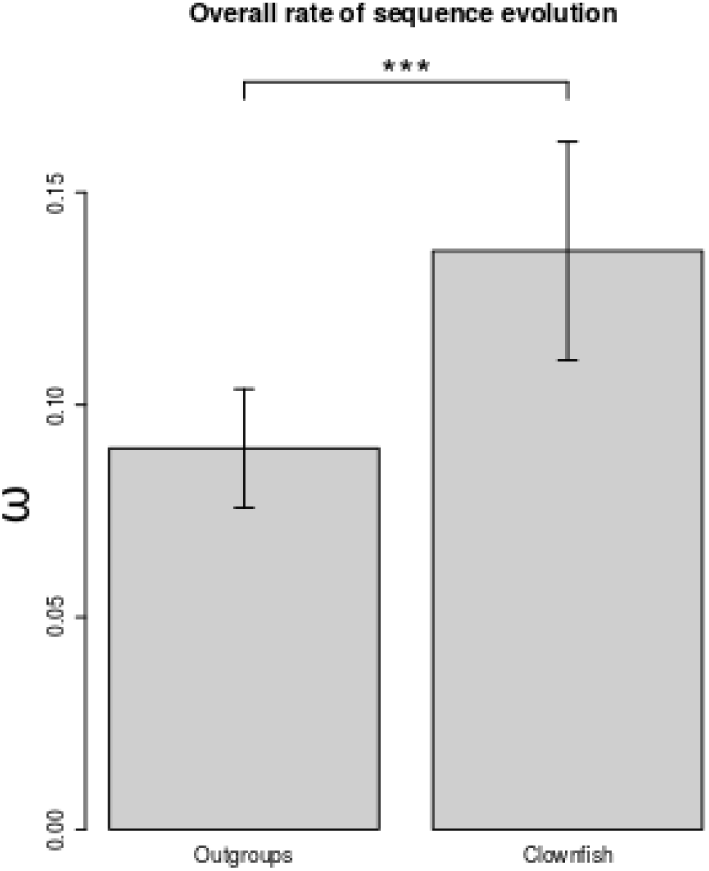
Overall rate of sequence evolution in outgroup species and clownfishes. The values of ω for outgroups and clownfishes were estimated with the branch model implemented in PAML, on 50 replicates of the concatenated alignments of 20 randomly picked genes. The significance of the difference in ω was tested with a two-sample Student’s t-test.

After correcting for multiple testing, we found a total of 732 genes that showed a significant signal of positive selection in the whole clade (5.4 % of the genes tested). The positively selected genes were distributed across the 24 chromosomes (between 21 and 50 significant genes per chromosome; Supplementary Figure S8). After normalizing by the total number of genes mapped to each chromosome, the percentage of positively selected genes was homogeneous across the chromosomes (Supplementary Figure S8). The proportion of sites under positive selection and the estimated ω for those sites varied across positively selected genes (Supplementary Figure S9). On average, the percentage of sites with signatures of positive selection (site class 2a and 2b, Figure 1) was 2.7%, with an average foreground ω of 19.2.

Positive selection analyses on 200 randomly selected genes using either the gene trees or the species tree resulted in similar results. Indeed, after correcting for multiple testing, we detected three concordant positively selected genes independently of the tree used (Supplementary Table S7). The data simulated under neutral or purifying selection scenarios showed that positive selection analyses resulted in the detection of no false-positives (Supplementary Figure S10). The power to detect positive selection increased with the increasing strength of selection (i.e., increasing ω, Supplementary Figure S10). While the power was only 5 % when we simulated an ω of 2, it increased to 54 %, 85 %, and 97.5 % for ω of 5, 10, and 20, respectively (Supplementary Figure S10).

Out of the 732 positively selected genes, 721 were functionally annotated. We found among the genes with the strongest signal of selection (Supplementary Table S8) those encoding for the isotocin receptor (ITR, OG20045_2b; Table 1) and the somatostatin 2 (SSTS, OG14285; Table 1). GO enrichment analysis performed on the positively selected genes resulted in 30 enriched GO terms (Supplementary Table S9). We found GOs linked to sexual reproduction (GO:0019953), detection of abiotic stimulus (GO:0009582), cellular response to interferon-gamma (GO:0071346), and cuticle development (GO:0042335). Within the positively selected genes participating in the enrichment of these GO terms (Supplementary Table S10), we found the genes encoding for the neuropeptide FF receptor 2 (NPFFR2, OG16291), the dual oxidase protein (DUOX, OG8036), and the rhodopsin (RHO, OG8544; Table 1).

### Parallel evolution associated with host and habitat divergence

Genes involved in the ecological divergence of clownfishes might evolve at different rates or show different selective pressures in specialists and generalists species. Thus, we investigated the presence of genes showing concerted differences in the evolutionary rate and selective pressures between all specialists and generalists species, which would suggest some level of parallel evolution in clownfish diversification.

We identified genes with a higher evolutionary rate in generalists or specialists in each species pair (Figure 7A and Supplementary Figure S11) and investigated whether these genes were shared among some of the species pairs (Figure 7B-C). The number of genes shared between 2, 3, 4, and 5 species pairs was similar in specialists and generalists (two-sample Student’s t.tests, *p-*values > 0.05, Supplementary Table S11; Figure 7B-C), and it quickly decreased with the increasing number of considered pairs (Figure 7B-C and Supplementary Figure S11). We did not detect any gene showing a higher evolutionary rate in all generalists (Figure 7B) or in all specialists (Figure 7C). When considering genes shared by 2, 3, or 4 species pairs, similar results were obtained for each combination of species pairs, independently from their phylogenetic relationship (Supplementary Figure S12).

**Figure 7:**
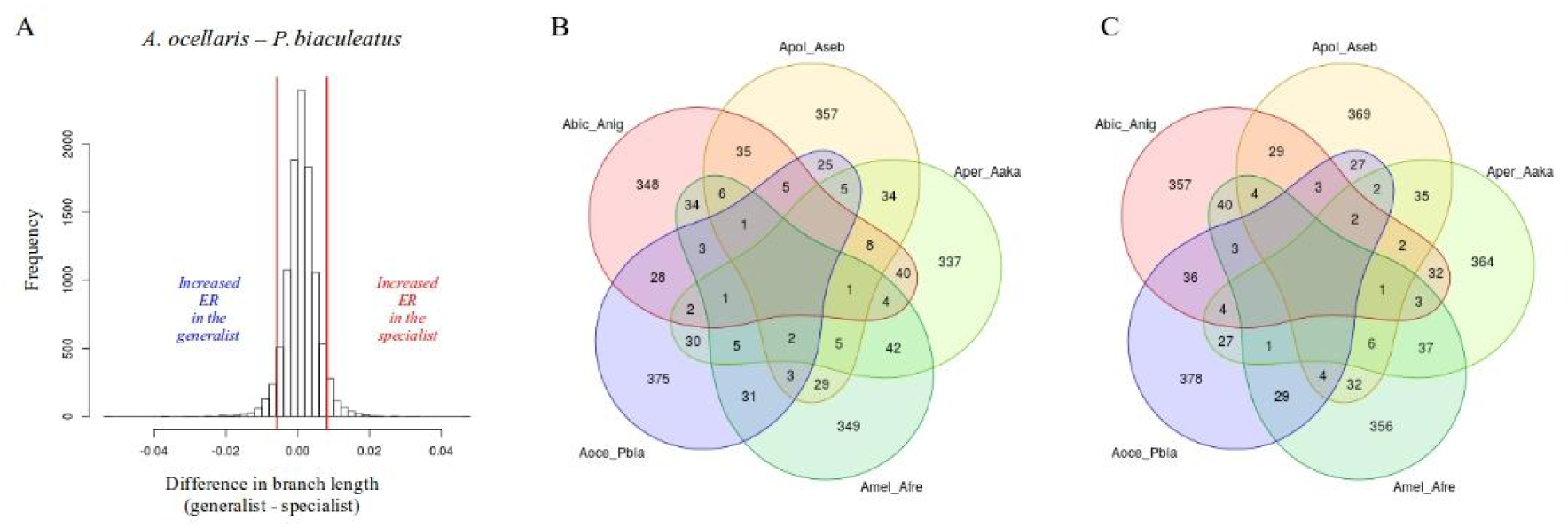
Evolutionary rate linked with host and habitat usage. A) Distribution of the branch length difference between the generalist *A. ocellaris* and the specialist *P. biaculeatus* obtained from the gene tree reconstructions for 11,273 OGs. The branch length was considered as a proxy for the evolutionary rate. We defined genes with a higher evolutionary rate in specialists and generalists those genes in the lower and upper 5% of the distribution, respectively. ER means evolutionary rate. The distributions of the additional species pairs are reported in Supplementary Figure S11). B) Number of genes with a higher evolutionary rate in generalists shared among the five species pairs. C) Number of genes with a higher evolutionary rate in specialists shared among the five species pairs. The species pairs correspond to: Abic_Anig: *A. bicinctus* and *A. nigripes;* Apol_Aseb: *A. polymnus* and *A. sebae*; Aper_Aaka: *A. perideraion* and *A. akallopisos*; Amel_Afre: *A. melanopus* and *A. frenatus*; Aoce_Pbia: *A. ocellaris* and *P. biaculeatus*.

We detected 3,991 genes that showed significantly different ω for specialists, generalists, and outgroups (clade model C better than null model M2a_rel; *p-*value corrected for multiple testing; Figure 1B). The vast majority of these genes (3,889 or 97.4%) presented signatures of purifying selection or neutral evolution (ω ≤ 1) in all specialists, generalists, and background branches. However, there were twice as many genes where the ω_specialists_ was estimated to be at least ten times lower than the ω_generalists_ (1,929 genes) than the reverse (955 genes). When exploring in more detail the genes with differences in ω between specialist and generalist species (Figure 2), we found a total of 155 genes showing relaxation or intensification of purifying selection in either specialists or generalists (Table 2). GO enrichment analysis of the genes showing intensified purifying selection in specialists resulted in 8 enriched GOs, including terms associated with the reproductive process (GO:0001541, ovarian follicle development; GO:0007283, spermatogenesis; Supplementary Table S12). Similarly, the enrichment of one GO term was found for genes with patterns of intensified purifying selection in generalists (GO:0046676, negative regulation of insulin secretion; Supplementary Table S12).

**Table 2:** Number of genes showing patterns of selection linked with host and habitat divergence. Information on purifying selected genes is available in Supplementary Table S13, while positively-selected genes are reported in Supplementary Table S14. The distribution of the ω values for each category is reported in Supplementary Figure S13 (for purifying selection) and Supplementary Figure S14 (for positive selection).

|  | Specialists | Generalists |
| --- | --- | --- |
| Relaxation from purifying selection | 4 | 16 |
| Intensification of purifying selection | 82 | 53 |
| Positive selection | 26 | 39 |

Genes with patterns of positive selection specific to specialists (ω_specialists_ > 1.5; ω_generalists_ and ω_background_ <= 1) and generalists (ω_generalists_ > 1.5; ω_specialists_ and ω_background_ <=1) were also detected (Table 2, Supplementary Table S14 and Supplementary Figure S14). GO enrichment analysis on genes positively selected in specialists resulted in two enriched GO terms: GO:0031398 (positive regulation of protein ubiquitination) and GO:0071456 (cellular response to hypoxia; Supplementary Table S12). Similarly, two enriched GO terms were detected for genes positively selected in generalists: GO:1903076 (regulation of protein localization to the plasma membrane) and GO:0006906 (vesicle fusion; Supplementary Table 12).

## DISCUSSION

In this study, we took advantage of the genomic data available for five pairs of closely related but ecologically divergent clownfish species to first investigate the genomic architecture of clownfish adaptive radiation and, second, explore if the evolution of their convergent phenotypes resulted from parallel genomic changes. Similarly to what was observed in other lineages undergoing adaptive radiation (e.g., Jones et al., 2012; Brawand et al., 2014; Fan & Meyer, 2014; Feiner, 2016; Berner & Salzburger, 2015; Faber-Hammond et al., 2019; Xiong et al., 2021), we found that clownfish genomes show bursts of transposable elements, overall accelerated rates of evolution of protein coding genes and topology inconsistencies potentially resulting from hybridization events. These characteristics possibly facilitated the rapid diversification of the group by generating genomic variations on which natural selection can act. Additionally, we observed that 5.4% of the genes showed a signature of positive selection throughout clownfish radiation. Among these genes, five have functions associated with social behavior and ecology (discussed below), and they could have played a role in the evolution of the size-based hierarchical social structure of clownfishes. Finally, we found genes with patterns of relaxation or intensification of purifying selection and genes with signals of positive selection linked with the ecological divergence, suggesting some level of parallel evolution during the diversification of the group.

### The overall genomic architecture of clownfish adaptive radiation

#### Burst in transposable element

The proportion of transposable elements (TEs) present in clownfishes is high (20-25% of the genome), and is comparable to what was observed in East African cichlids (Brawand et al., 2014; Shao et al., 2019) and in *A. percula* genome (Lehmann et al., 2019). This result confirms a reliable annotation of the TEs despite the higher fragmentation of the assemblies used in this study. This large TEs proportion in clownfishes originated mainly from two bursts of transpositions. The older burst happened in the ancestor of *O. niloticus* and the Pomacentridae, and it was also observed in the cichlids (Shao et al., 2019) and *P. moluccensis* (Supplementary Figure S6). The more recent burst occurred around the split of *P. moluccensis* from the common ancestor of clownfishes. This increase in transpositions was also observed in *P. moluccensis* but was less intense and rapidly decreased (Supplementary Figure S6), suggesting that the burst started in the *P. moluccensis-*clownfishes ancestor and, after the split of the species, only continued in clownfishes.

Bursts in TEs transpositions are common in fishes and likely favor species diversity (Shao et al., 2019). Indeed, the movement of TEs can lead to insertions, deletions, and chromosomal rearrangements (Lönnig & Saedler, 2002; Fedoroff et al., 2012), potentially contributing to adaptation, speciation, and diversification processes (Syvanen, 1984; Rebollo et al., 2010, Auvinet et al., 2018). The high percentage of TEs observed in clownfishes, and the increased burst of transposition compared to *P. moluccensis* might therefore have facilitated the diversification of the group, as observed in other adaptive radiations such as the East African cichlid fishes (Brawand et al., 2014; Fan & Meyer, 2014), *Heliconius* butterflies (Dasmahapatra et al., 2012; Lavoie et al., 2013) and *Anolis* lizards (Feiner, 2016).

We did not detect any main difference in the TEs content or TEs transposition histories for the different clownfish species. Nevertheless, species-specific genomic features may be underestimated here as the assemblies were based on the *A. frenatus* reference genome (Marcionetti et al., 2019).

#### Accelerated coding evolution

We detected a significantly increased evolutionary rate in a subset of randomly sampled clownfish genes, suggesting a global accelerated coding evolution in the group. Interestingly, accelerated morphological evolution in clownfish compared to damselfish has also been previously reported (Litsios et al., 2012). Accelerated evolution in genes involved in morphological and developmental processes was observed in Eastern African cichlids (Brawand et al., 2014) and potentially facilitated their rapid diversification (Brawand et al., 2014; Berner & Salzburger, 2015). Because accelerated coding evolution was widespread in clownfish, pinpointing specific biological processes with higher evolutionary rates and potentially involved in the diversification (as performed in Brawand et al., 2014) has not been possible.

Overall differences in evolutionary rates between lineages are widely observed (e.g., Ohta, 1993; Bromham et al., 1996; Bromham, 2002; Nabholz et al., 2008; Bromham, 2009; Welch et al., 2008; Thomas et al., 2010) and are primarily attributed to differences in generation-time, where organisms with shorter generation times likely evolve faster (i.e., the generation-time hypothesis, Bromham, 2009). However, the generation time in clownfishes (5 years estimated on *A. percula*, Buston & García, 2007) is higher than in the considered outgroup species (i.e., from a few months to 4 years, Supplementary Table S15), excluding it as the driver of the acceleration in coding evolution. The effective population also affects the evolutionary rate through the action of genetic drift (Lynch & Walsh, 2007), and lineages with smaller effective population sizes (*N*_*e*_) show an increased rate of evolution (Woolfit & Bromham, 2003, 2005). Although *N*_*e*_ for the species in the study was not available, sequential hermaphroditism can potentially reduce it (Chopelet et al., 2009; Coscia et al., 2016; Benvenuto et al., 2017; but not in Waples et al., 2018), and clownfishes are the only sex-changing species considered here (data from FishBase; Froese & Pauly, 2000). Thus, further studies investigating *N*_*e*_ are needed to evaluate its effect on the observed accelerated coding evolution of clownfishes.

Coevolution between species can also alter the rate of molecular evolution. Antagonistically coevolving species are expected to show an increased rate of molecular evolution (the Red Queen Hypothesis, Van Valen, 1971; Pal et al., 2007; Paterson et al., 2010), while species in mutualistic relationships should show a decrease in their rate of evolution (the Red King Effect theory, Bergstrom & Lachmann, 2003). Nevertheless, similarly to what detected here for clownfishes, an increased evolutionary rate was observed in obligate symbiotic organisms (Lutzoni & Pagel, 1997; Yoshizawa et al., 2003; Bromham et al., 2013) and mutualistic ants (Rubin & Moreau, 2016). Demographic history (e.g., *N*_*e*_ and generation time) could explain the increased rate in symbiotic organisms, but not in the mutualistic ants (Rubin & Moreau, 2016). In this less intimate mutualistic interaction, species must adapt to their own changing environments and those of their symbionts, which may lead to selective pressure similar to those experienced by antagonistically coevolving species (Van Valen, 1971) and lead to an increased rate of evolution (Rubin & Moreau, 2016). This hypothesis could also hold for the clownfish mutualism, especially given the presence of generalist species living in different hosts (see also below). Thus, while we cannot exclude an effect of demographic history without further analyses, the increased evolutionary rate observed in clownfishes may also result from the acquisition of mutualism with sea anemones.

#### Positive selection in clownfishes

We found that at least 5.4% of the genes were positively selected throughout clownfish diversification. This percentage could be underestimated, as simulations showed a low power in detecting weak selection and demonstrated the absence of false positives. Positive selection on ecologically important genes can drive the adaptation and diversification of organisms (e.g., Rundle & Nosil, 2005; Schluter & Conte, 2009), and patterns of positive selection were observed in the genomes of radiating lineages (e.g., Kapralov et al., 2013; Brawand et al., 2014; Cornetti et al., 2015; Nevado et al., 2016). While linking all positively selected genes with a potential role in clownfish diversification would be unreasonable, we identified five genes with particularly interesting functions that we discuss further.

First, we detected positive selection on the gene encoding the somatostatin 2 (SST2, OG14285, Table 1). Somatostatins are a diverse family of peptide hormones that influence organismal growth by inhibiting the production and release of the growth hormone (Very & Sheridan, 2002; Sheridan & Hagemeister, 2010). The variation in somatic growth rate resulting from social status changes in the fish *Astatotilapia burtoni* was associated with a shift in the volume of somatostatin-containing neurons (Hofmann & Fernald, 2000). Similarly, in *A. burtoni*, somatostatin 1 regulates aggressive behavior in dominant males (Trainor & Hofmann, 2006), and its expression in the hypothalamus is associated with the control of somatic growth depending on social status (Trainor & Hofmann, 2007). Second, we observed positive selection on the gene encoding the isotocin receptor (ITR, OG20045_2b, Table 1). In the cichlid *Neolamprologus pulcher*, isotocin plays an important role in modulating social behavior, increasing responsiveness to social information (Reddon et al., 2012, 2015, 2017). For instance, exogenous administration of isotocin resulted in increased submissive behavior when challenged aggressively (Reddon et al., 2012), and the size and number of isotocin neurons were significantly different between cooperatively and independently breeding species (Reddon et al., 2017). We also detected positive selection on the gene *NPFFR2* (OG16291, Table 1), encoding for the neuropeptide FF receptor 2, which plays a role in feeding-related processes (including appetite control, food intake, and gastrointestinal motility) in *Lateolabrax maculatus* (Li et al., 2019).

Within sea anemones, clownfishes are organized in a size-based dominance hierarchy, with the female and the male being respectively the largest and the second-largest individuals. The non-breeders (if present) are progressively smaller as the hierarchy is descended (Fricke, 1979; Ochi, 1989). These well-defined size differences between individuals are maintained by the precise regulation of the subordinate growth (Buston, 2003), often through aggressive behavior (Iwata et al., 2008, 2010). It was also hypothesized that, in *A. percula*, subordinates reduce their food intake to avoid exceeding size thresholds, which could lead to conflicts with dominants (Chausson et al., 2018). Thus, positive selection in somatostatin 2, isotocin receptor and *NPFFR2* genes possibly contributed to the evolution of the size-based dominance hierarchy in clownfishes through the modulation of both growth and aggressive/submissive behaviors.

We also observed positive selection in the *RHO* gene (OG8544, Table 1), encoding the rhodopsin. This protein is a photoreceptor required for image-forming vision at a low light intensity, and modifications in its gene sequence likely lead to changes in the absorbed wavelength (Bowmaker, 2008). In cichlids, positive selection on rhodopsin was associated with shits in the wavelength absorbance of fish at different depths, promoting ecological divergence in the group (Spady et al., 2005; Sugawara et al., 2009). Divergent positive selection on this gene was also observed among lake and river cichlid species (Schott et al., 2014). Clownfishes live at depths between 1 and 40 meters, with the depth depending on the habitat of their host sea anemones (Fautin & Allen, 1997). Although we did not detect explicit positively-selected sites affecting rhodopsin absorbance, this gene may have played a role in clownfish adaptation to different depths.

Finally, we detected positive selection in the *duox* gene (OG8036, Table 1), encoding the dual oxidase protein involved in synthesizing thyroid hormones (Chopra et al., 2019), which regulates the white stripes formation in *A. percula* (Salis et al., 2021). They also observed that shifts in *duox* expression and thyroid hormone levels due to ecological differences resulted in the divergent formation of stripes and color patterns in *A. percula* (Salis et al., 2021). The functional role of striped patterns in clownfish is still unknown but could be associated with the sea anemone ecology (Salis et al., 2021) or with species recognition, as observed in other teleosts (Seehausen et al., 1999; Kelley et al., 2013). Positive selection on *duox* in clownfishes may thus be associated with divergence in the formation of white stripes in the group.

While these genes show interesting functions associated with clownfish social behavior and ecology, further investigation is needed to validate the link between the detected positively selected genes and their role in clownfish evolution and diversification. For instance, expression evidence for these genes is lacking (except for *duox*) and should be examined in the future.

#### Mosaic genomes in clownfishes

The reconstruction of the mitochondrial and nuclear phylogenetic trees of clownfishes resulted in cytonuclear discordance. Nodes in both the mitochondrial and nuclear phylogenetic trees were strongly supported, suggesting that the inconsistency results from past hybridization events or incomplete lineage sorting (ILS). Cytonuclear incongruences in clownfish were previously detected and were associated with a burst in the diversification rate of the group (Litsios & Salamin, 2014). We also observed topological discordance throughout the nuclear genome, with phylogenetic trees inferred in the different genomic regions being well supported, confirming potential hybridization events and/or ILS in the clownfish diversification. The main topological inconsistencies were seen in the deep nodes of the trees, while the five pairs of closely related species mainly branched together. A first exception was detected for a few windows, in which the specialists *A. nigripes* and *A. sebae* branched as sister species, which we will further discuss below. We observed a second exception for *P. biaculeatus*, which principally branched as sister species of *A. ocellaris* but was also frequently placed as basal to the *Amphiprion* clade (Figure 4C). This disparity mirrors the conflicting phylogenies reported in the literature (e.g., Frédérich et al., 2013, Mirande, 2017; na Ayudhaya et al., 2017, Lobato et al., 2014; DiBattista et al., 2016). While *Premnas* has been recently recovered within *Amphiprion* (Tang et al., 2021), these inconsistencies suggest a complex evolutionary history of *P. biaculeatus* - such as potential hybridization events with species outside the *Amphiprion* genus - that should be further investigated.

As the topological inconsistencies were observed in the deep nodes of the trees, hybridization events and/or ILS likely happened in ancestral clownfish species. Following the topology of the nuclear phylogenetic tree, the second most abundant topology was the one of cluster 1 (i.e., not the one reflecting the mitochondrial tree). This topology was predominant in chromosomes 1 and 3, suggesting the presence of mechanisms (such as structural inversions, decreased recombination, selection, and/or genetic drift) increasing its fixation (see also below). However, to identify such mechanisms and characterize the topological landscape of clownfish more accurately, the precise identification of topological breakpoints directly on the *A. percula* reference genome is necessary.

We observed another particular topological inconsistency on chromosome 18, where the most frequent topology split the species pair of *A. perideraion - A. akallopisos* (topology of cluster 5, Figure 4C) and suggested past gene flow between *A. perideraion* and the *A. melanopus* - *A. frenatus ancestor*. This topology was almost exclusively observed on this chromosome and clustered in two large regions, potentially indicating that the introgression signal was removed from the rest of the genome by extensive backcrossing but persisted on chromosome 18 through the disruption of recombination. The recombination disruption was possibly achieved through the genomic inversions of the regions - originated in the ancestor of *A. melanopus* - *A. frenatus* and introgressed in *A. perideraion*, or arisen in *A. perideraion* after the gene flow - that were fixed in *A. perideraion* through genetic drift or selection. Genomic inversions that break recombination, creating clusters of loci controlling ecologically important traits, consequently fixed by natural selection, are observed in the case of supergenes (e.g., Joron et al., 2011; Kunte et al., 2014; Zinzow-Kramer et al., 2015; Küpper et al., 2016; Branco et al., 2018). Here, however, we cannot establish a role of selection, as we cannot easily link the genes’ functions in these regions to species’ important ecological traits without further studies (Supplementary Table S4), and we did not observe an enrichment of positively selected genes on chromosome 18. Thus, we cannot exclude the role of genetic drift in fixing these regions in *A. perideraion*. It is worth mentioning that the evolution of sex chromosomes may also result in particular patterns as those observed on chromosome 18 (e.g., Natri et al., 2019). However, we do not believe that this is relevant in clownfishes, as these species are sequential hermaphrodites with no sex chromosomes (Fricke & Fricke, 1977; Moyer & Nakazono, 1978; Fricke, 1979; Arai, 2011), and genes involved in the sex change are scattered throughout the genome (Casas et al., 2018).

The patterns observed on chromosome 18, together with the additional topological inconsistencies, suggest ancestral hybridization events in the diversification of clownfishes. Gene flow spreading ancient genetic variation among species has been proposed to facilitate adaptive radiation (Berner & Salzburger, 2015; Marques et al., 2019). For instance, ancestral hybridization between distinct lineages has fueled the adaptive radiation of cichlids (i.e., the *hybrid swarm* hypothesis, Seehausen, 2004; Meier et al., 2017; Svardal et al., 2020), while introgressive hybridization among members of the radiating lineages (i.e., the *syngameon* hypothesis, Seehausen, 2004) has facilitated ecological speciation in *Heliconius* butterflies (Dasmahapatra et al., 2012; Pardo-Diaz et al., 2012) and Darwin’s finches (Lamichhaney et al., 2015). While hybridization events may have also participated in clownfish diversification, we cannot yet exclude that the mosaic genomes observed in clownfishes are, at least partially, the result of ILS. Further studies should focus on the formal testing of gene flow, for instance, with ABBA-BABA tests and related Patterson’s *D* statistics (Green et al., 2010; Durand et al., 2011) as performed in Maier et al. (2017).

#### No increase in gene duplication rate at the basis of clownfish radiation

We observed the highest duplication rate in the common ancestor of the Pomacentridae and *O. niloticus* (Supplementary Figure S7), comparable to the estimated duplication rate in the cichlids’ ancestor (Figure 2 in Brawand et al., 2014). Contrarily, the duplication rate detected in clownfish was similar to the one observed in non-radiating teleosts (Figure 2 in Brawand et al., 2014). Thus, clownfish diversification does not seem to be characterized by an increase in gene duplication events, differently from what has been observed in cichlids adaptive radiation (Brawand et al., 2014; Machado et al., 2014). While the overall duplication rate may be underestimated in a phylogenetic duplication analysis approach (as performed here) compared to read depth or array comparative genomic hybridization (aCGH) methods, the relative difference in the rate between branches remains consistent among the analyses (see Brawand et al., 2014), which reinforce the validity of our findings.

Although we did not observe an increased gene duplication rate in clownfishes, we tested if the duplication events were, nevertheless, followed by positive selection in the whole group. Indeed, gene duplications allow for the divergent evolution of the resulting gene copies, permitting functional innovation of the proteins and/or expression patterns (Lynch & Conery, 2000; Taylor & Raes, 2004; Kondrashov, 2012). We did not detect any duplicated gene positively selected in all clownfishes and thus associated with the diversification of the whole clade. It is worth mentioning that the method we employed here tests for signals of positive selection in all branches and may result in a reduced power due to multiple testing (Smith et al., 2015). Nevertheless, it was important to perform the analysis in an exploratory way, as we did not know beforehand which copy of the genes was potentially positively selected. Anyhow, if some duplicated genes were indeed positively selected in all clownfishes, the intensity of positive selection was not strong enough to permit their detection.

### Parallel evolution in clownfish diversification

In clownfish, we did not detect extensive topological inconsistencies potentially arising from hybridization events among specialist or generalist species (Figure 4C), indicating that parallel evolution through recent introgressive hybridization is not a major driver of clownfish diversification. Nevertheless, we observed an exception in a few genomic windows, where the two specialists *A. nigripes* and *A. sebae* branched as sister species. The node of this split was fully supported in all windows. The species *A. sebae* is part of the primary radiation of clownfishes, which diversified in the Coral Triangle region (Litsios et al., 2014), and this species experienced range expansion after the diversification (Litsios et al., 2014) to reach its current distribution in the Northern Indian Ocean (including Sri Lanka and Maldives; Allen, 1991, Litsios et al., 2014). In contrast, *A. nigripes* is endemic to the Maldives and Sri Lanka (Allen, 1991, Litsios et al., 2014) and belongs to the replicated radiation of clownfishes, which diversified independently in the Indian Ocean after a colonization event (Litsios et al., 2014). While there is no report of sympatric populations of the two species, their geographical distribution suggests that they co-occur in the Maldives and Sri Lanka, and gene flow between them is thus conceivable.

If introgression between the species occurred, extensive backcrossing likely removed its signal from the rest of the genome, and natural selection possibly fixed the regions showing topological inconsistency. Genes within these regions included olfactory receptors and the *Has2* gene, encoding for hyaluronan synthase 2, which catalyzes the addition of GlcNac to nascent hyaluronan polymers (Tian et al., 2013). Interestingly, it is well known that coral reef fish larvae use olfactory cues to identify suitable settlement sites (e.g., Lecchini et al., 2005a,b; Gerlach et al., 2007; Dixson et al., 2008). Additionally, genes with functions associated with GlcNAc and hyaluronan were previously found positively selected at the basis of clownfish radiation and were hypothesized to play a role in the acquisition of mutualism by avoiding the release of sea anemones toxins. (Marcionetti et al., 2019). These genes might be involved in the ecological convergence of *A. nigripes* and *A. sandaracinos*, with introgressive hybridization playing a role in the parallel evolution of species, as observed in *Heliconius* butterflies (Dasmahapatra et al., 2012; Supple et al., 2013). Nevertheless, future studies that aim at better characterizing the regions of topological inconsistency (using *A. percula* reference genome), better evaluating the functions and expression of the genes, and better defining the phenotypes involved, are needed before drawing any conclusion.

While parallel evolution through recent introgressive hybridization does not seem frequent in clownfish diversification, parallel evolution could also result from differential rates of evolution or selective pressures in genes associated with ecological divergence (e.g., Cresko et al., 2004; Colosimo et al., 2005; Linnen et al., 2013; Projecto-Garcia et al., 2013). Regarding the evolutionary rate, we did not detect any gene evolving at different rates in all the species pairs. It is nevertheless worth noting that, when considering only some of the species pairs, we observed genes with an increased evolutionary rate shared among specialists or generalists (Figure 7B-C). However, the number of shared genes rapidly decreased with the increasing number of species pairs considered. The decrease was similar in specialists and generalists and was independent of the phylogenetic relationship between species pairs. These observations suggest that the shared genes were likely obtained by random processes rather than being actually related to ecological divergence.

Contrary to the evolutionary rate, we observed genes undergoing similar selective pressures in all specialist or generalist clownfish species. Indeed, we detected 24 and 39 genes that were specifically positively selected in specialists and generalists, respectively. Because they show patterns of positive selection, these genes were potentially involved in the parallel adaptation to similar ecological niches, which resulted in the convergent morphological changes observed in clownfishes (Litsios et al., 2012). Genes with parallel patterns of relaxation or intensification of purifying selection in specialists or generalists species were also observed. The selective pressures on these genes may reflect parallel outcomes of the adaptation to similar ecological niches. These results suggest the presence of some level of parallel evolution during the diversification of clownfishes. Nevertheless, in general, a clear link between the function of these genes and their potential role in - or how they are affected by - clownfish adaptation to similar ecological niches cannot be drawn without a better characterization of clownfish functional traits.

It is worth mentioning that, in general, we detected stronger purifying selection in specialist species compared to generalists. This observation is in accord with theoretical expectations postulating that, overall, generalist species experience relaxed selection because they use multiple environments, leading to a decrease in the efficiency of selection (Kawecki, 1994; Bono et al., 2020; Draghi, 2021). Additionally, by considering the rationale behind the increased evolutionary rate reported for mutualistic ants (Rubin & Moreau, 2016), generalist species might show a higher rate of sequence evolution as their environments are more changing than those of specialists (i.e., going in the direction of the Red Queen Hypothesis, Van Valen, 1971).

## CONCLUSIONS

Studies on well-described adaptive radiations have started investigating the intrinsic genomic factors that may promote the rapid diversification of lineages. Here, we examined the genomic features underlying clownfish diversification.

Similar to other adaptive radiations, our results showed that clownfish genomes show two bursts of transposable elements, overall accelerated coding evolution, and topology inconsistencies potentially resulting from hybridization events. These characteristics possibly promoted clownfish adaptive radiation. Additionally, we detected positively selected genes with interesting functions, which likely participated in the evolution of the size-based dominance hierarchy in the group. Finally, we observed genes that underwent differential selective pressures associated with ecological divergence, suggesting some extent of parallel evolution during the group’s diversification.

This study provides cues for the further investigation of the genomics underlying clownfish adaptive radiation. For instance, along with the necessity to better characterize clownfish’s functional traits, future studies will need to formally test for ancestral hybridization events in the clownfish diversification. Additionally, other genomic features proposed to have facilitated rapid diversification - such as regulatory element changes, structural variants or heterozygosity - should also be investigated. Estimating clownfishes’ effective population sizes is also essential to understand the mechanisms underlying the observed acceleration in coding sequence evolution in the group. Nevertheless, our results provide the first genomic insights into clownfish adaptive radiation and integrate the growing collection of studies investigating the genomic mechanisms governing species diversification.

## ACKNOWLEDGEMENTS

We thank B. Micheli and S. Schmid for general support. We also thank the Vital-IT and the DCSR infrastructure of the University of Lausanne. Funding: University of Lausanne funds, Swiss National Science Foundation, Grant Number: 31003A-163428

## Notes

### Competing Interest Statement

The authors have declared no competing interest.

