## Supplementary Information and Figures for "Insights into the genomics of clownfish adaptive radiation: the genomic substrate of the diversification"

### Table of content

Supplementary Information

- S1. Gene trees vs species tree for selection and evolutionary rate analyses Page 3
- S2. Gene duplication analysis Page 4

Supplementary Tables – Legends Page 6

Supplementary Figures Page 8

- **Figure S1.** Bootstrap support for the trees reconstructed along the genomes. Distribution of the average bootstrap values for all windows (left) and distribution of node support for all nodes and windows combined (right).
- **Figure S2.** Summary of all the tree topologies reconstructed along the genome. The summary was obtained with DensiTree.
- **Figure S3.** DensiTree summary of topology of cluster 6 of *treespace*. The cluster contained many different topology, and could not be summarized by a single topology.
- **Figure S4.** Alternative topology found in 13 windows of the genome. The two specialists species *A. nigripes* and *A. sebae* branching as sister species.
- **Figure S5.** Overall proportion of TEs (left) and proportion of each major TE class (right) in clownfish, *P. moluccensis* and additional Actinopterygii genomes. Clownfishes and *P. moluccensis* genomes were annotated with RepeatMasker (http://www.repeatmasker.org/). Information on TE content for the additional species was retrieved from previous studies (Brawand et al., 2014; Gao et al., 2106; Lehmann et al., 2019).
- **Figure S6.** Kimura distance-based copy divergence analysis of transposable elements in the clownfishes and *P. moluccensis*.
- **Figure S7.** Divergence calculated as the neutral genomic divergence between clownfishes and six outgroup species and duplication rates in four internal branches (on the right).

Results for duplication rate are reported for the internal branches labelled as A, B, C and D, and reported in the tree. Branch A corresponds to the ancestral branch of clownfishes. The duplication rate was calculated as the number of duplication events in the branch normalized by the branch length.

- **Figure S8.** Number of positive selected genes in each of the clownfishes chromosomes. Red line correspond to the proportion of genes that are positively selected in each chromosome.
- **Figure S9.** Proportion of sites under positive selection and the estimated ω for those sites
- **Figure S10.** False positive and power in detecting positive selection in the whole clownfish clade.
- **Figure S11.** Difference in evolutionary rate for the four species pairs. The missing pairs is reported in Figure 7A.
- **Figure S12.** Number of genes with higher evolutionary rate in generalists (A, C) and specialists (B, D) species. In A and B, we reported the number of genes when considering the genes present in the reported species. In C and D we reported the number of genes when the reported species is missing.
- **Figure S13.** Omega values for specialists, generalists and background for genes showing intensification or relaxation from purifying selection in generalists and specialists.
- **Figure S14 .** Omega values for genes positively-selected in specialists and generalists
- **Figure S15.** Duplication events observed in each branch of clownfishes and outgroups. The duplication numbers on each branch are reported in red. The branch length are reported in black.

### Supplementary Information

#### S1. Gene trees vs species tree for selection and evolutionary rate analyses

The tree topology affects the inference of selection (Diekmann and Pereira-Leal, 2015) and can bias the analyses of the evolutionary rate associated with host usage. When topological incongruence exists, the use of either gene trees or the species tree may lead to inconsistent results. The species topology does not always represent the correct gene genealogy across the whole genome, while the gene tree reconstruction can result in inaccurate topologies due to sampling errors (Planet, 2006). This is particularly the case in highly-conserved protein-coding genes, where the amount of variable sites is limited. Additionally, topological accuracy in gene trees also decreases with alignment errors (Ogden & Rosenberg, 2006).

We performed Shimodaira-Hasegawa tests (SH-test; Shimodaira and Hasegawa, 1999) between the species tree and all gene trees to assess the level of topological incongruence. This test assesses whether a set of selected trees are equally good explanations of the data (null hypothesis) or if a set of trees are significantly better (alternative hypothesis). We computed the likelihood of the species and gene trees given the alignments with the *pml* and *optim.pml* functions implemented in the phangorn R package (v.2.4.0; Schliep, 2011), and we performed the SH-test with the *SH.test* function of the same library. We classified the OGs into three categories: “better gene trees” (GT; 206 OGs), “better species trees” (ST; 2,021 OGs), and “non-significant” (NS; 11,273 OGs). We verified that genes in the GT category were not located in genomic regions showing alternative topologies (see *Mosaic genomes in clownfishes*). We visually inspected these alignments and computed the following alignment statistics: alignment length, number of gaps, and number of variable sites. As OGs in this category showed overall shorter and less accurate alignments (Supplementary Table S16), their gene trees probably resulted from alignment errors, and we did not consider them further.

The analysis of the evolutionary rate was performed on the gene trees, but only for genes in the NS category. We did not consider OGs in the ST category, as using gene trees for these OGs would not have been accurate (i.e., gene trees are not a good explanation of the data). We performed the positive selection analyses using the species tree considering the OGs in the NS and ST categories. We nevertheless verified the effect of using either the species or gene trees (Diekmann and Pereira-Leal, 2015) by randomly selecting 200 1-to-1 OGs and repeating the positive selection analyses following the same procedures (*see Positive selection analyses on the whole clownfish group*) but using the gene tree. We compared the results by visually investigating the log-likelihood obtained for each model. We examined whether the same OGs were inferred as positively selected after multiple-testing corrections of the *p-values*, and we verified that similar estimates of ω were obtained with the species and gene trees.

### S2. Gene duplication analyses

#### Methods

We investigated the gene duplication events occurring during the diversification of clownfishes and the other fish species *P. moluccensis*, *S. partitus, O. niloticus*, *G. aculeatus*, and *T. nigroviridis*. We retrieved 2,725 multicopy HOGs and filtered out the gene copies of the additional outgroup species not considered here. We employed a phylogenetic duplication analysis (PDA) approach (similar to Brawand et al., 2014), counting the number of gene duplication events observed at each branch of the phylogenetic trees of these species. Within each HOG, a gene duplication event was counted when the number of gene copies in all the species descending from that branch was higher than the maximum number of gene copies in all other species. This approach was conservative, as it did not consider parallel gene duplications (i.e., duplications events happening independently in different taxa) or gene losses. However, it reduced the errors in the deeper branches caused by misassembled genes. Indeed, fragmented gene annotations or separate assemblies of divergent alleles may be mistaken for paralogous genes in terminal branches. This bias is decreased in internal branches as an increasing number of species must display the higher number of gene copies. Because of this, we only considered duplication events in the internal branches.

The number of duplication events in each branch was normalized to account for the divergence between species. We estimated the neutral genomic divergence between the species using ca. 7.5 million fourfold degenerate sites that we obtained from the codon alignments of 1-to-1 OGs. We extracted the fourfold degenerate sites in each alignment and we concatenated them into a single alignment before reconstructing the phylogenetic tree with RaxML (GTR+Γ model, 100 bootstraps; v.8.2.12; Stamatakis, 2014). The number of duplication events detected on each branch was then divided by the corresponding branch length to obtain the duplication rate (i.e., the number of duplications normalized by the neutral divergence between the species).

#### Results

We explored whether clownfishes showed an increased rate of gene duplication that could be associated with their diversification. We found a total of 1,747 HOGs with specific duplication events across the full phylogenetic tree (Supplementary Figure S7 and S15). We detected 19 duplication events in the common ancestor of clownfishes (Supplementary Figure S7 and S15). In contrast, we observed 90 duplications both in the common ancestor of the Pomacentridae (Supplementary Figure S7 and S15) and before the split of *P. moluccensis* and the clownfishes (Supplementary Figure S7 and S15). When normalizing for species divergence, the duplication rates were similar, with about 12 duplicated genes/percent of divergence (Supplementary Figure S7). A higher duplication rate was observed in the common ancestor of the Pomacentridae and *O. niloticus* (40 duplications/percent of divergence; Supplementary Figure S7).

We performed positive selection analyses on the duplicated genes (multicopy HOGs; see main text). We analyzed 2,725 HOGs and, while we found 116 genes with signals of positive selection in clownfish species, selection was only acting on a few clownfish branches. Thus, we did not detect positive selection throughout the whole clade potentially associated with the group's diversification (schematic in Figure 1A). Similarly, we did not observe any HOG with signals of positive selection linked with host and habitat use and thus associated with the evolution of convergent phenotypes (schematic in Figure 1B).

### Supplementary Tables - Legend

- **Table S1:** The 10 selected clownfish species and their host sea anemones. Data obtained from Fautin & Allen, 1997; Litsios et al., 2012.
- **Table S2:** Reference sequences used for the mitochondrial reconstruction of the clownfish species and *P. moluccensis.*
- **Table S3:** Statistics for the whole genome alignments of the ten clownfish species and *P. moluccensis.* Results for the alignments are based on *A. frenatus* reference. The number of scaffolds and the total assemblies length were obtained by averaging on the 11 species.
- **Table S4:** Significant Gene Ontology (GO) terms obtained for the regions of alternative topology on chromosome 18.
- **Table S5:** Locations of topologies with specialist species branching as sister species. Gene content and function of the windows with these topologies are also reported.
- **Table S6:** Percent of transposable elements content in clownfishes and *P. moluccensis* genomes.
- **Table S7:** Results for the positive selection analysis on the whole clownfish clade, using either the species tree or the gene trees. The analysis was performed on 200 randomly selected 1-to-1 OGs.
- **Table S8:** Subset of the positively-selected genes in the whole clownfish group. The genes falling in the upper 90% quantile of the foreground ω distribution are reported.
- **Table S9:** Gene ontology (GO) enrichment analysis on genes positively-selected in the whole clownfish clade. The analysis was performed with TopGO, using the fisher's exact test and the weight01 algorithms.
- **Table S10:** Annotation for positively-selected genes associated with enriched GO terms.
- **Table S11:** Shared number of genes in increasing species pairs for generalist and specialist species. No gene was found shared between 5 species in either generalists or specialists.
- **Table S12:** Gene ontology (GO) enrichment analysis on genes showing intensification of purifying selection in specialist clownfish species (A) and generalists (B), as well as genes showing patterns of positive selection in specialists (C) and generalists (D).
- **Table S13:** Genes showing selection patterns linked with host and habitat divergence.

- **Table S14**: Genes showing positive selection in specialists (A) and generalists (B) species.
- **Table S15:** Generation time of outgroup species used in the study.
- **Table S16:** Alignment statistics for the 1-to-1 OGs. Categories were defined from the results of SH-test, with GT, ST and NS categories corresponding to “better gene tree”, “better species tree” and “non-significant”, respectively. Because the genes in the GT category showed overall shorter and less accurate alignments, their trees probably resulted from alignment errors, and we did not consider them further.

**Figure S1. Bootstrap support for the trees reconstructed along the genomes.** Distribution of the average bootstrap values for all windows (left) and distribution of node support for all nodes and windows combined (right).

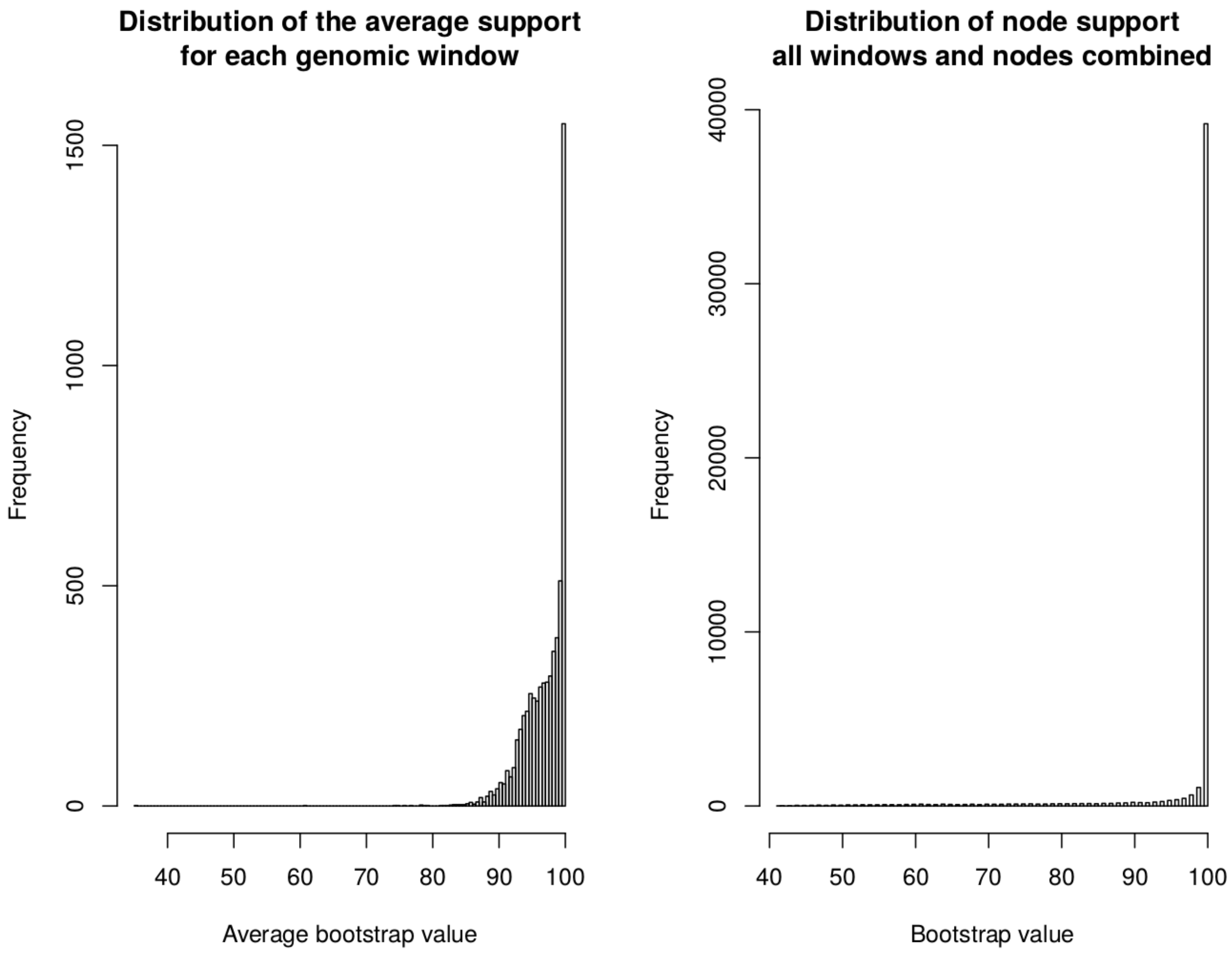

**Figure S2. Summary of all the tree topologies reconstructed along the genome.** The summary was obtained with DensiTree.

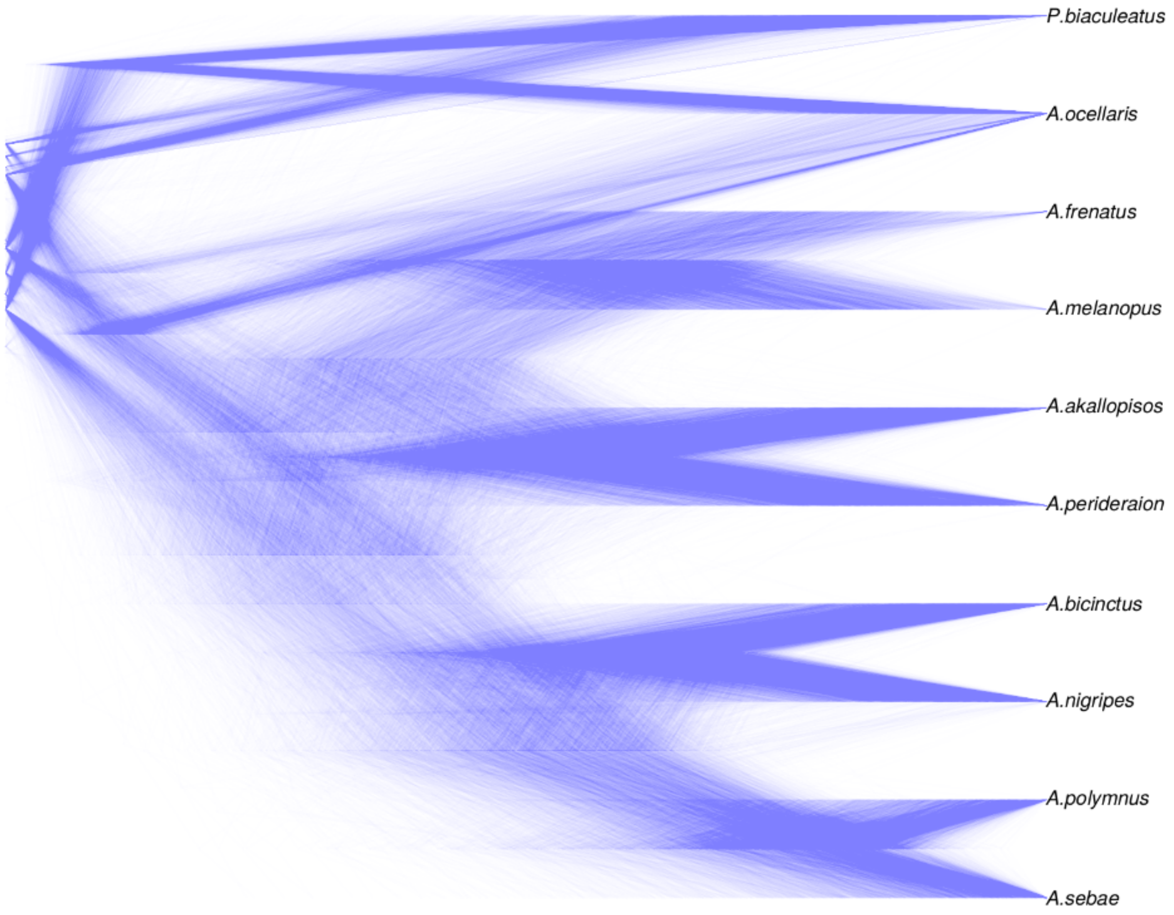

**Figure S3. DensiTree summary of topology of cluster 6 of *treespace*.** The cluster contained many different topology, and could not be summarized by a single topology.

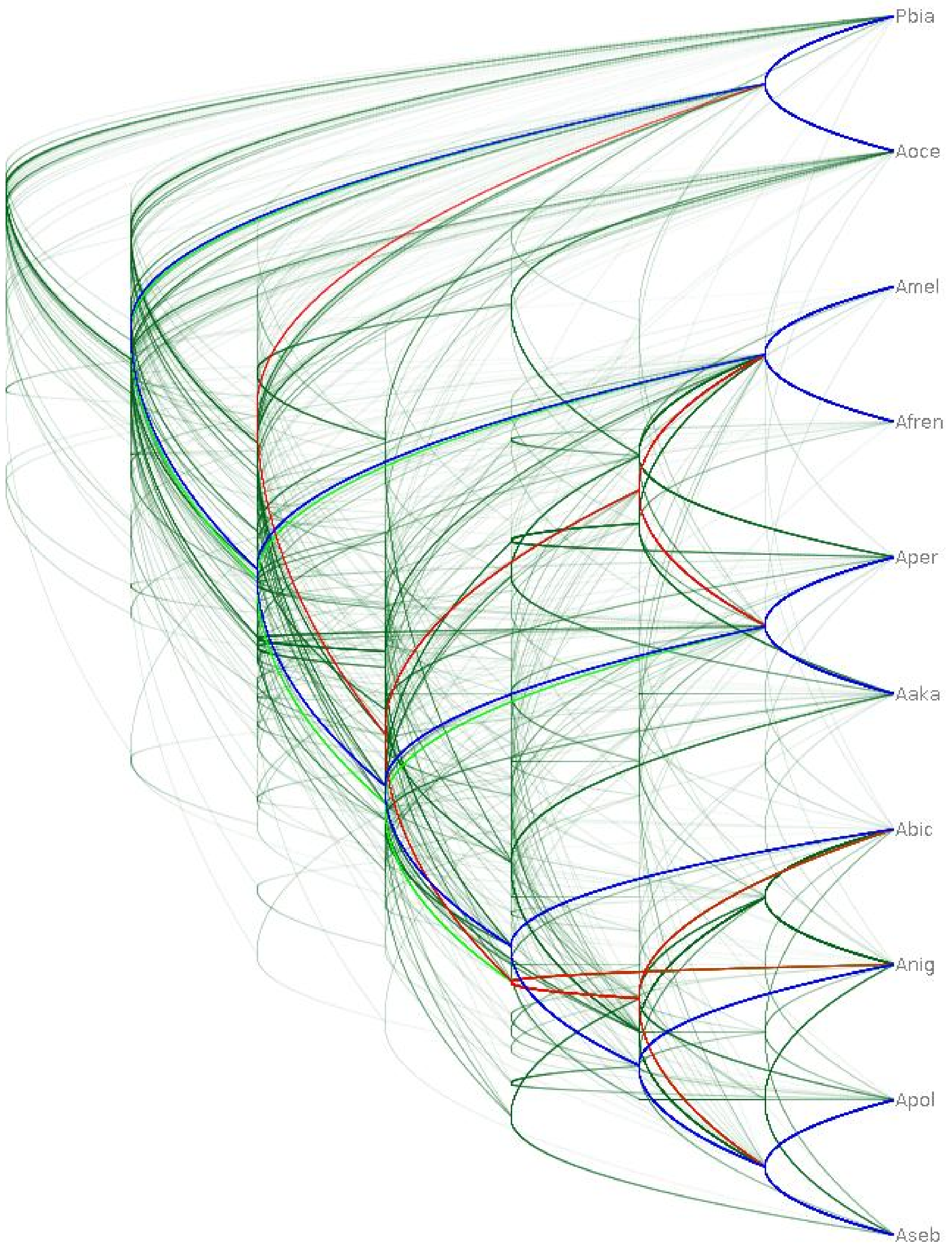

**Figure S4.** Alternative topology found in 13 windows of the genome. The two specialists species *A. nigripes* and *A. sebae* branching as sister species.

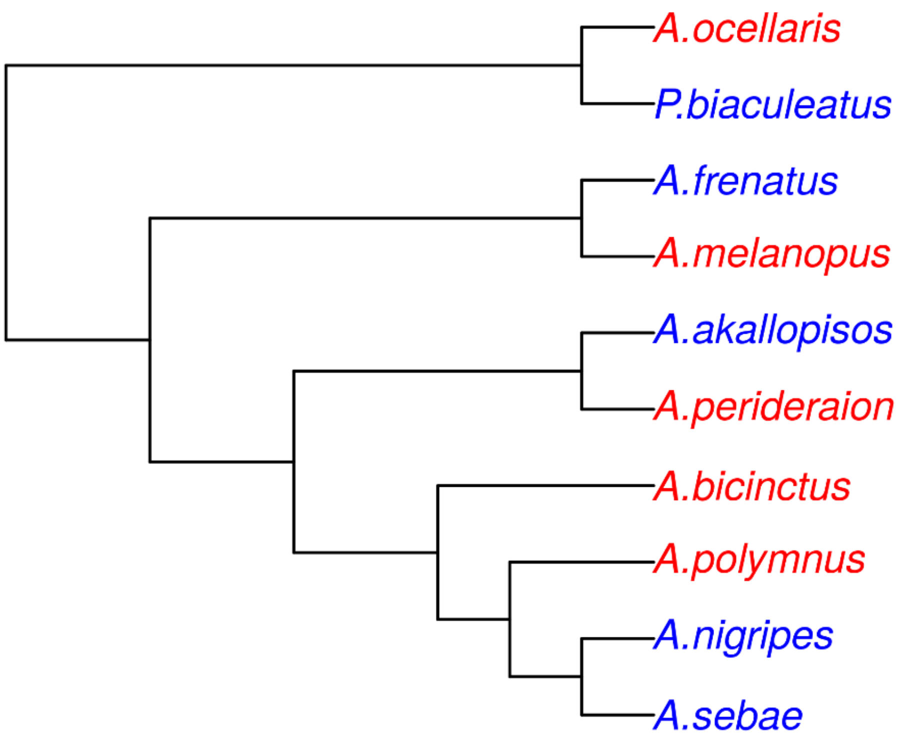

**Figure S5. Overall proportion of TEs (left) and proportion of each major TE class (right) in clownfish, *P. moluccensis* and additional Actinopterygii genomes.** Clownfishes and *P. moluccensis* genomes were annotated with RepeatMasker (http://www.repeatmasker.org/). Information on TE content for the additional species was retrieved from previous studies (Brawand et al., 2014; Gao et al., 2106; Lehmann et al., 2019).

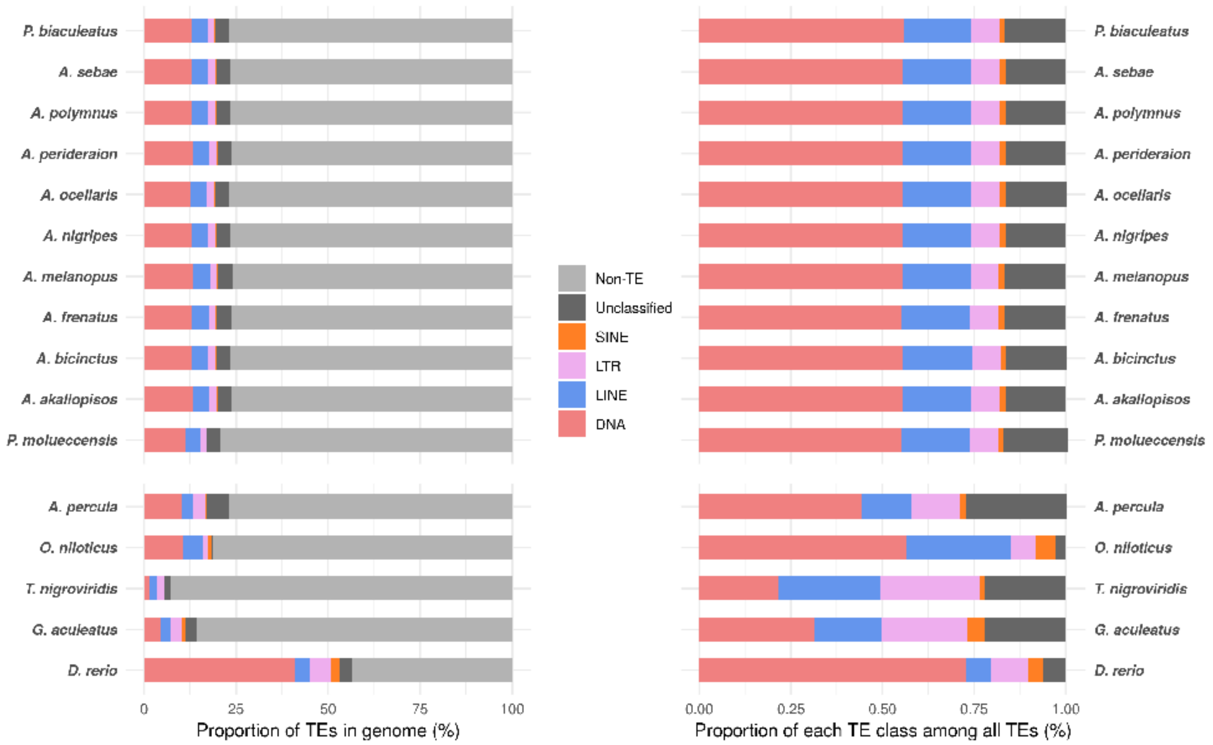

**Figure S6.** Kimura distance-based copy divergence analysis of transposable elements in the clownfishes and *P. moluccensis*.

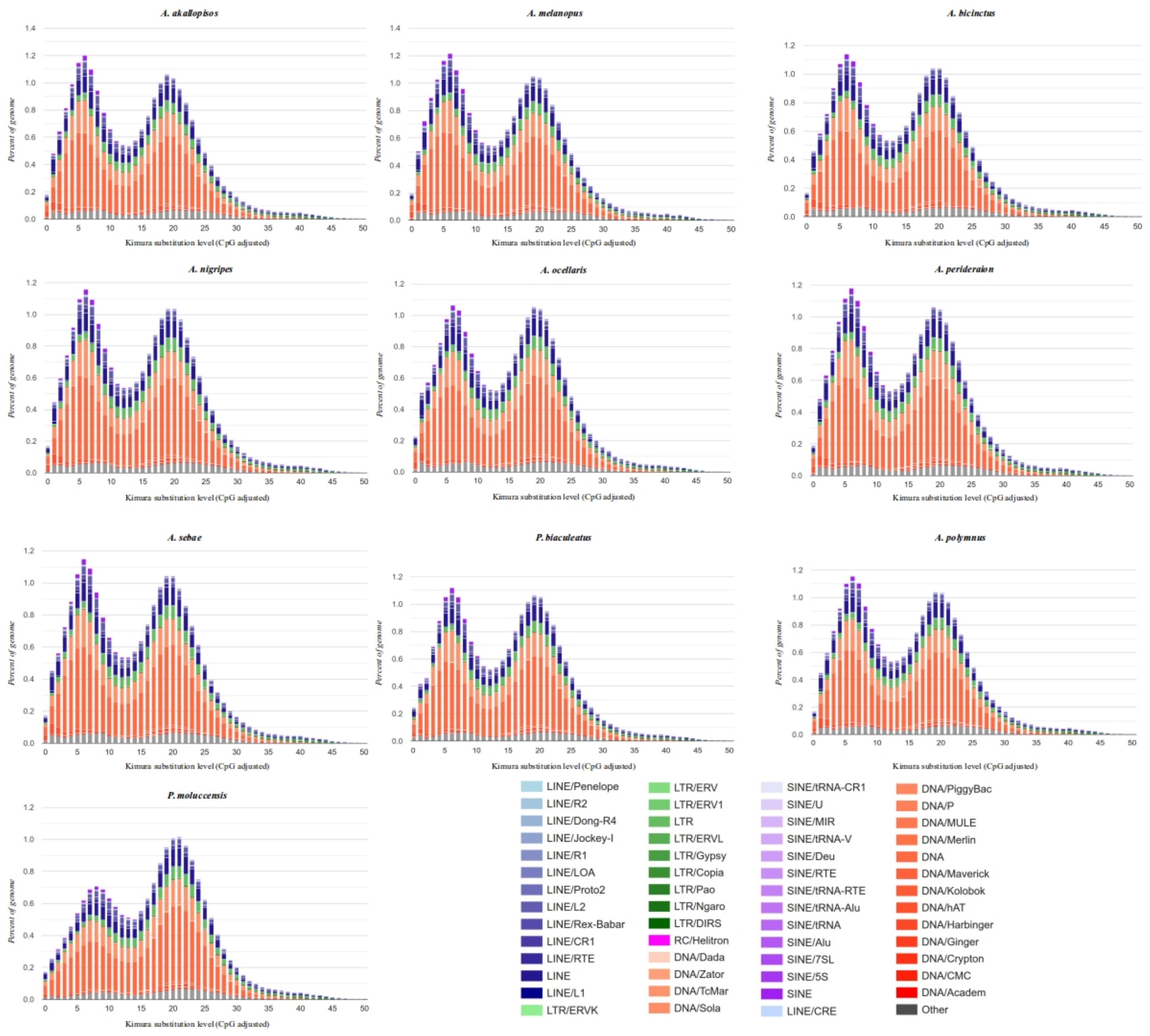

**Figure S7.** Divergence calculated as the neutral genomic divergence between clownfishes and six outgroup species and duplication rates in four internal branches (on the right). Results for duplication rate are reported for the internal branches labelled as A, B, C and D, and reported in the tree. Branch A corresponds to the ancestral branch of clownfishes. The duplication rate was calculated as the number of duplication events in the branch normalized by the branch length.

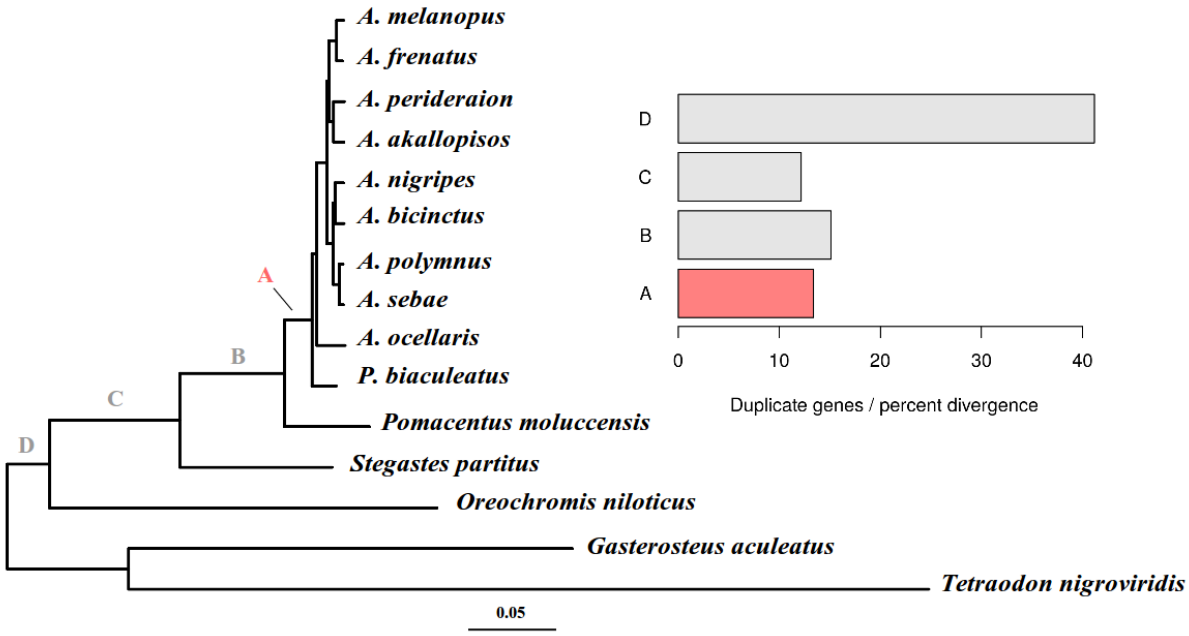

**Figure S8.** Number of positive selected genes in each of the clownfishes chromosomes. Red line correspond to the proportion of genes that are positively selected in each chromosome.

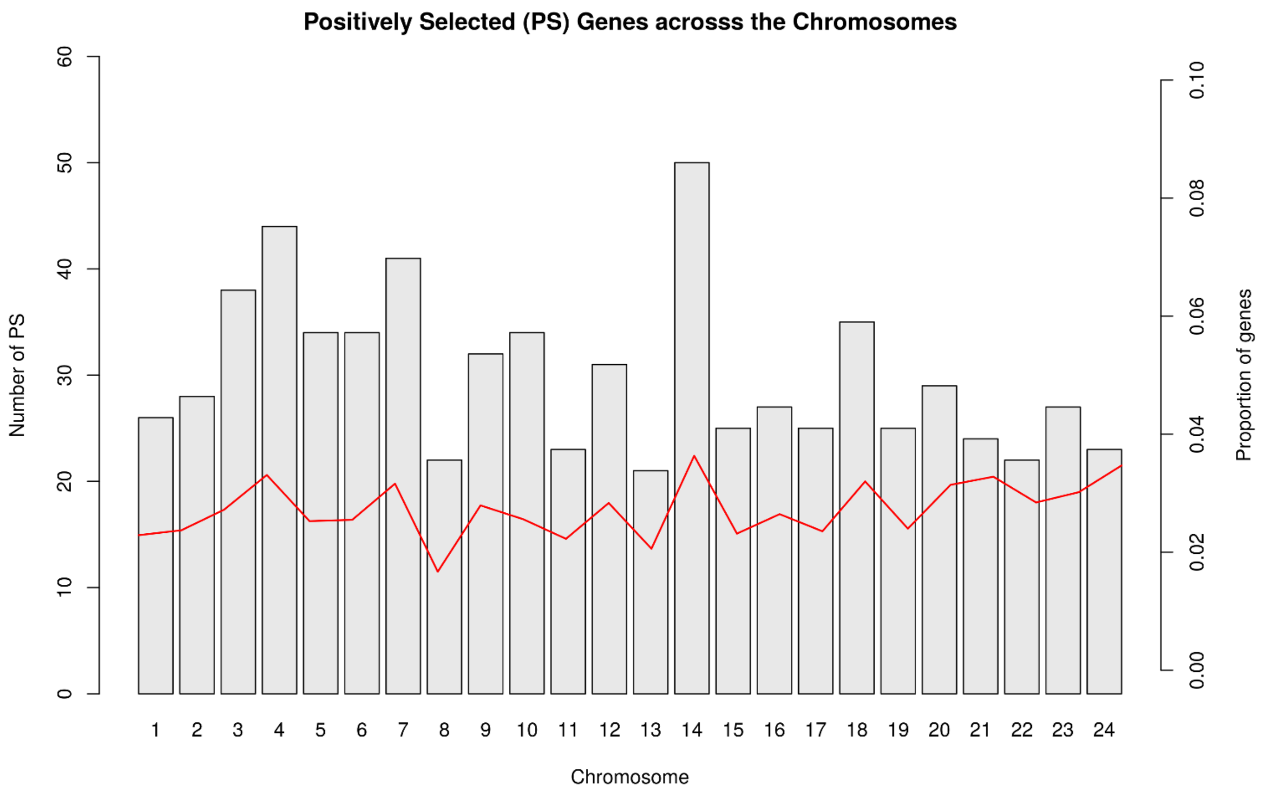

**Figure S9.** Proportion of sites under positive selection and the estimated ω for those sites.

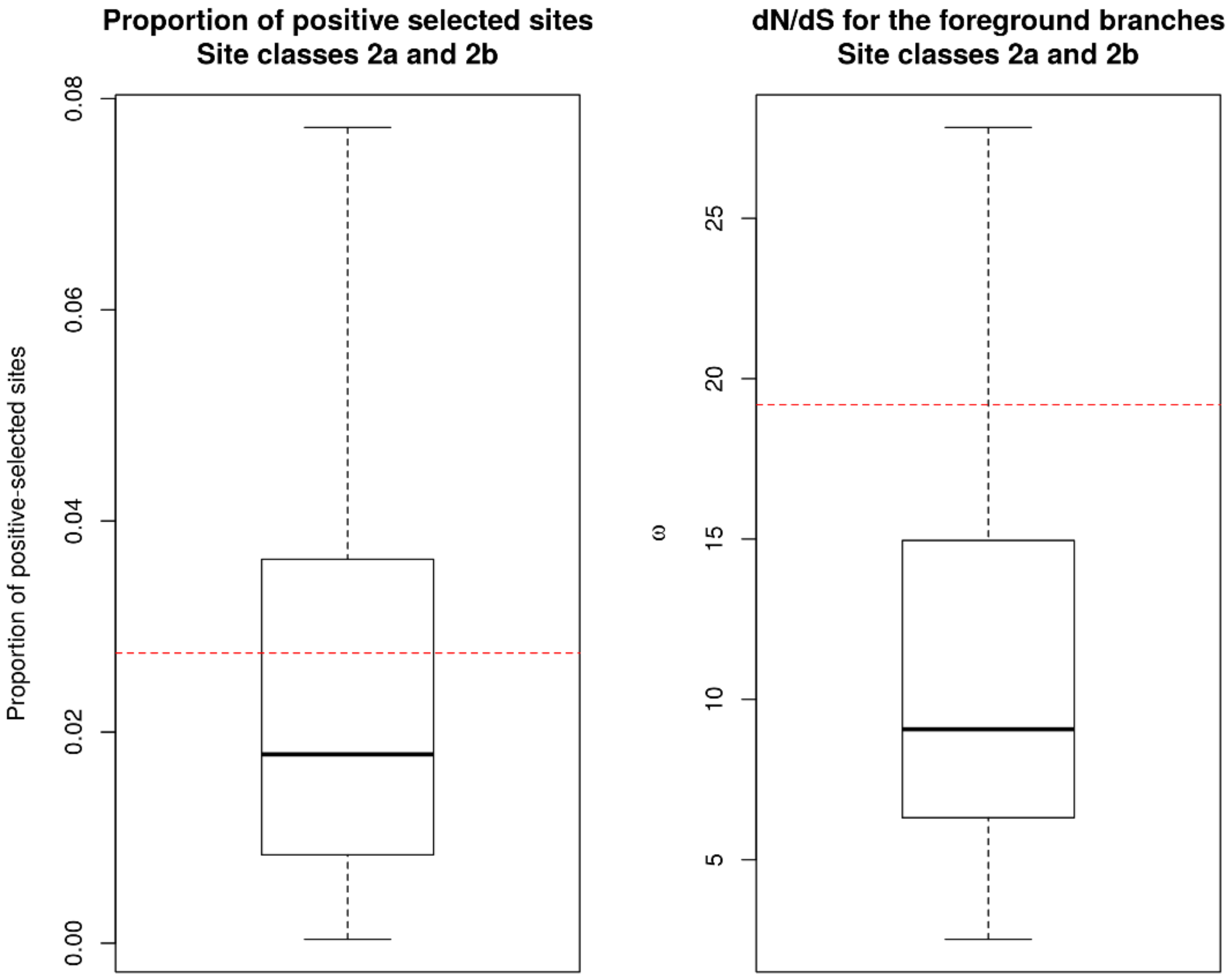

**Figure S10.** False positive and power in detecting positive selection in the whole clownfish clade.

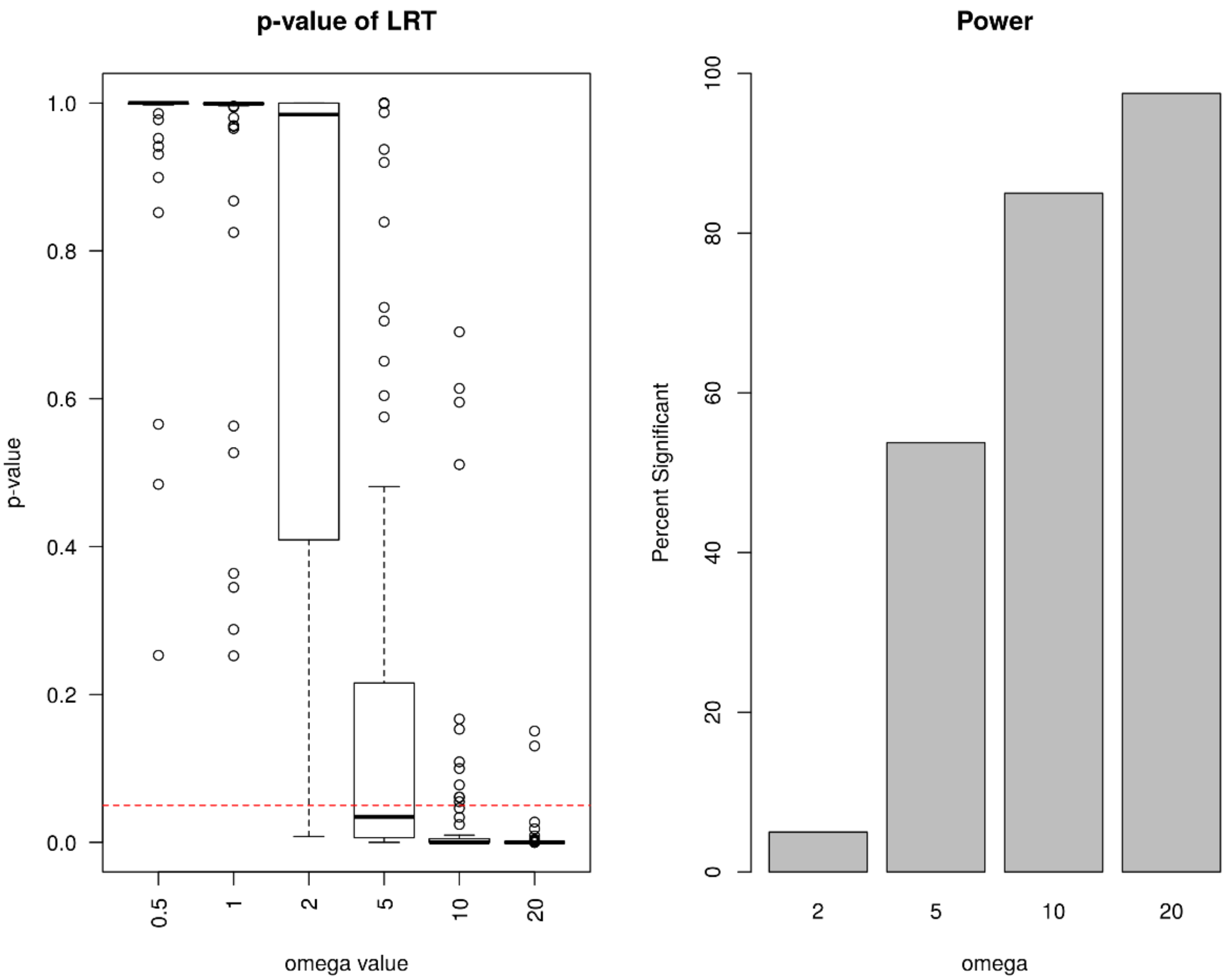

**Figure S11.** Difference in evolutionary rate for the four species pairs. The missing pairs is reported in Figure 7A.

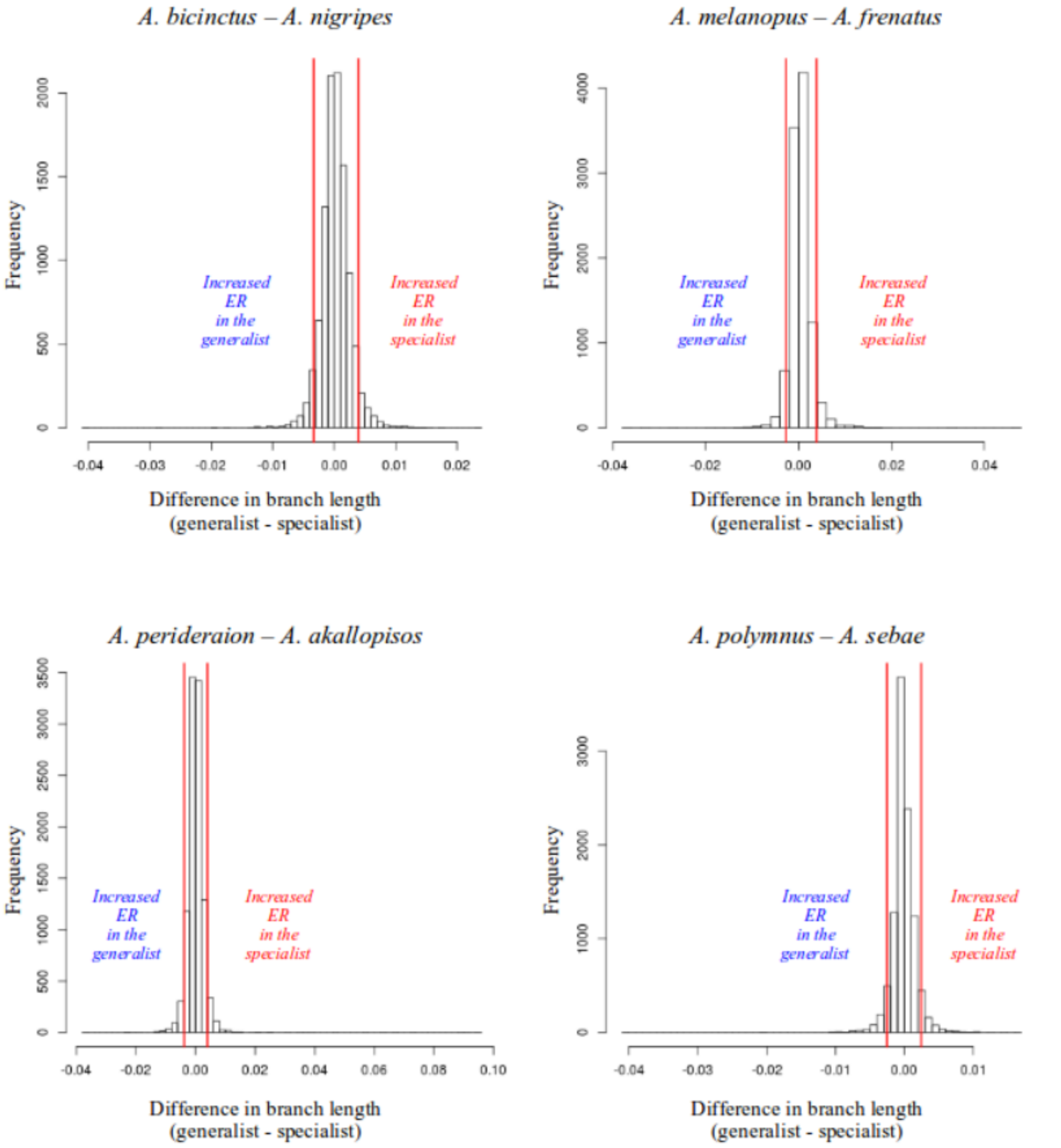

**Figure S12.** Number of genes with higher evolutionary rate in generalists (A, C) and specialists (B, D) species. In A and B, we reported the number of genes when considering the genes present in the reported species. In C and D we reported the number of genes when the reported species is missing.

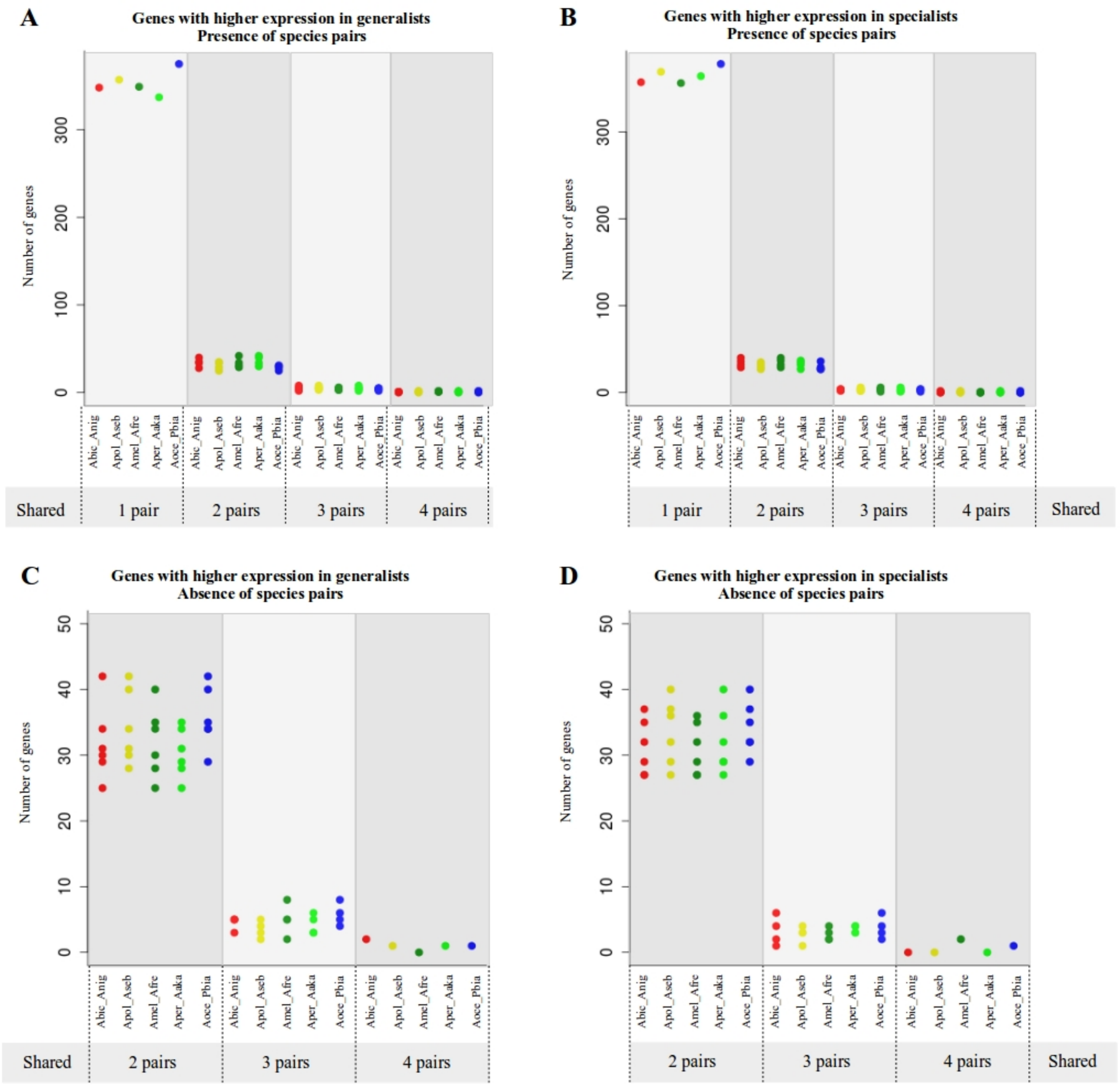

**Figure S13.** Omega values for specialists, generalists and background for genes showing intensification or relaxation from purifying selection in generalists and specialists.

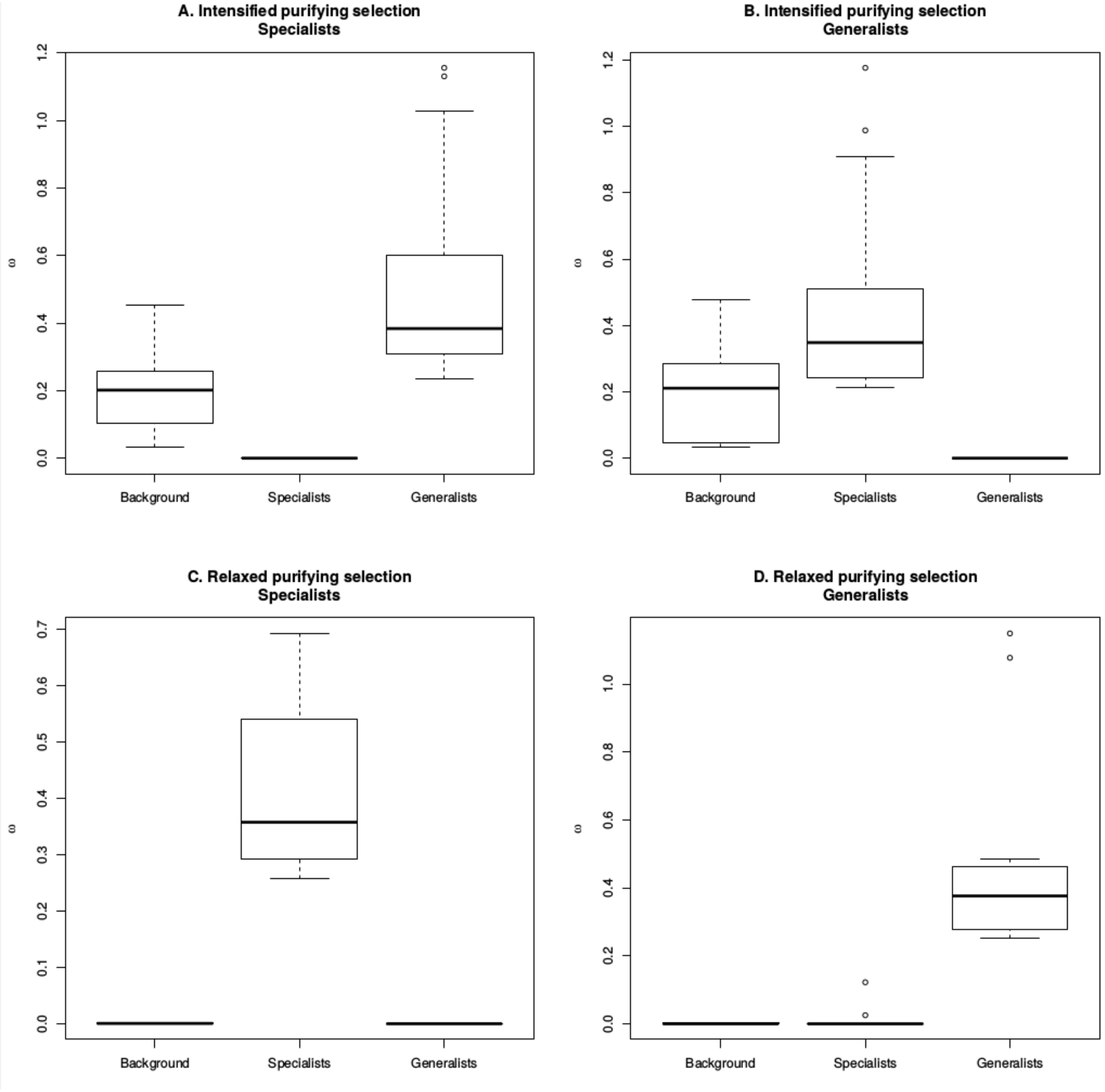

**Figure S14.** Omega values for genes positively-selected in specialists and generalists

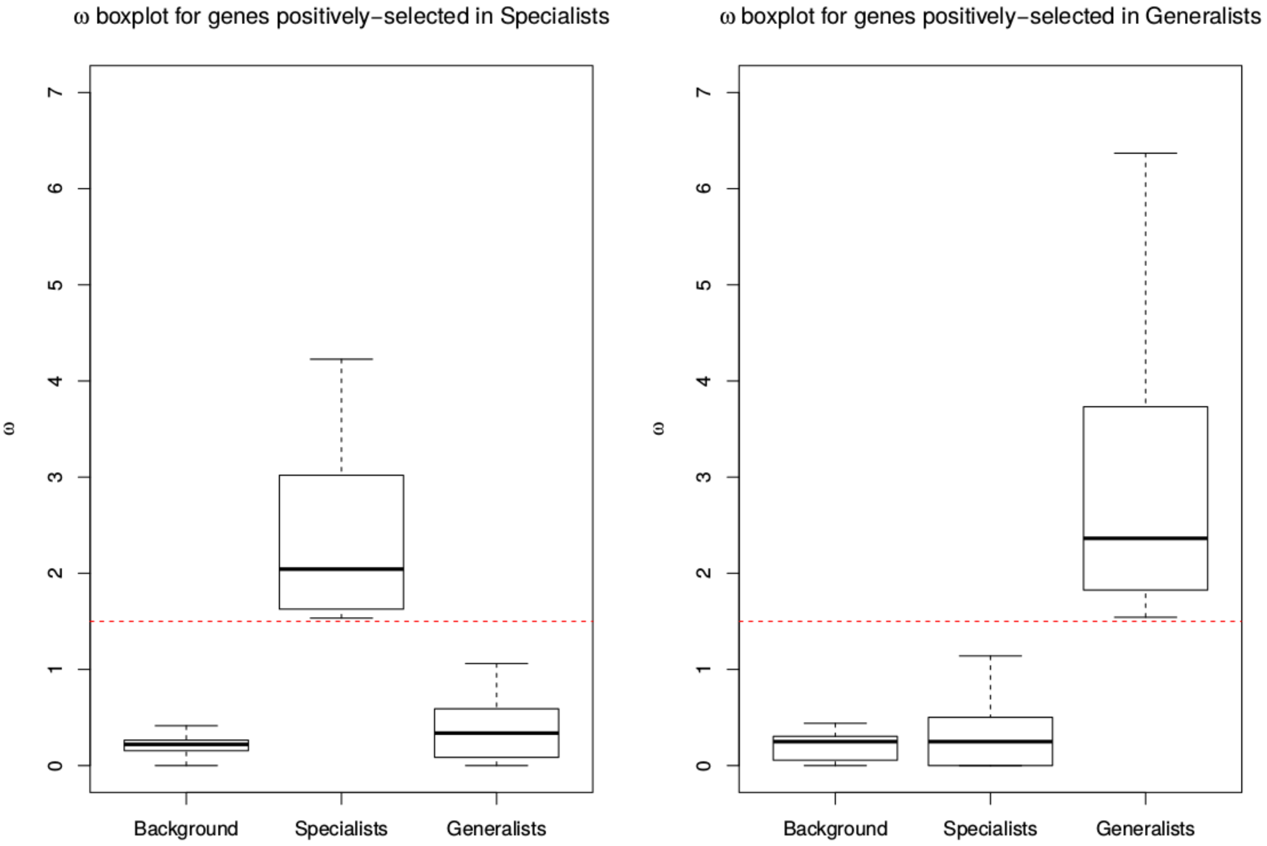

**Figure S15.** Duplication events observed in each branch of clownfishes and outgroups. The duplication numbers on each branch are reported in red. The branch length are reported in black.

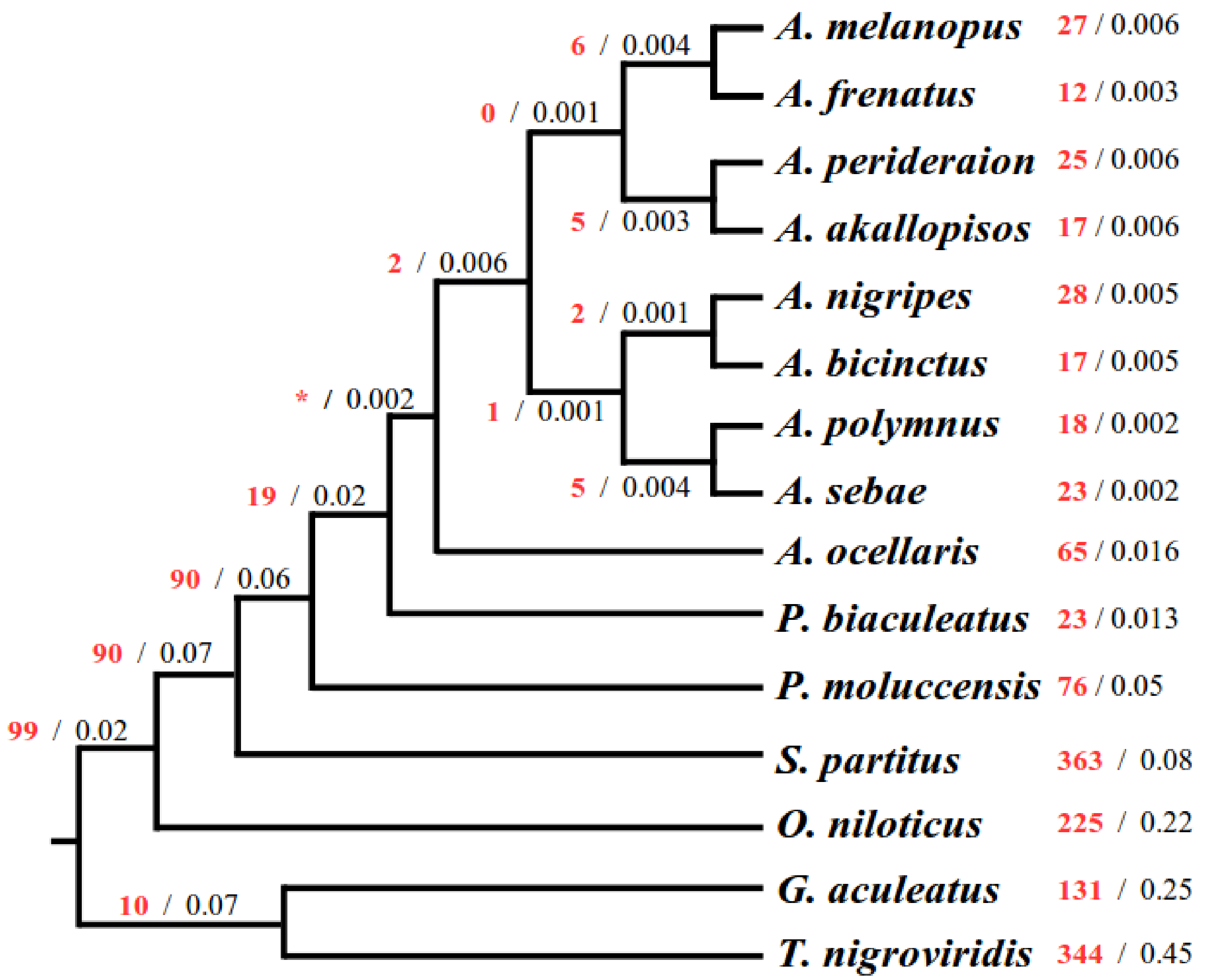
